# A Bayesian overhaul of thermal tolerance landscape models: Predicting ectotherm lethality buildup and survival amid heatwaves

**DOI:** 10.1101/2024.01.23.576827

**Authors:** Jahangir Vajedsamiei, Niklas Warlo, H. E. Markus Meier, Frank Melzner

## Abstract

1. In the face of escalating heatwaves, accurately forecasting ectotherm population mortality is a pressing ecological challenge. Current Thermal Tolerance Landscape (TTL) models, while surpassing single-threshold metrics by incorporating individual survival times, are constrained by frequentist regression parametrization reliant on constant-temperature experiments, omitting probabilistic outcomes.

2. This study addresses these limitations by pioneering the application of Approximate Bayesian Computation-Sequential Monte Carlo (ABC-SMC) to analyze survival data from Baltic *Mytilus* mussels subjected to both microcosm (constant temperature) and mesocosm (dynamic temperature) heatwave regimes.

3. The ABC-SMC yields probabilistic predictions of individual lethality buildup and population survival trajectories, closely aligned with observed survival data across both experimental conditions. Informed by more realistic dynamic data, the TTL model predicts local mussel resilience against the most extreme summer heatwaves projected for this century, albeit with considerations for sublethal impacts and potential recruitment declines.

4. Our approach can enhance the predictive accuracy concerning the sensitivity of key marine populations amidst intensifying heatwaves, addressing the urgent need for accurate modeling tools to inform conservation practices and ecosystem management, ultimately aiding in the preservation of marine biodiversity.

## INTRODUCTION

Anthropogenic emissions drive changes in global climate patterns, including ocean warming, acidification, and deoxygenation, with a notable rise in the frequency and intensity of extreme weather conditions, such as marine heatwaves (Laufkötter *et al*. 2020; IPCC 2021). Defined as extended episodes of anomalously elevated seawater temperatures, marine heatwaves become especially hazardous during summer when baseline temperatures are already high (Hobday *et al*. 2016; Holbrook *et al*. 2019; Laufkötter *et al*. 2020).

These anomalies pose acute threats to ectothermic species in shallow marine habitats, as their physiological functions are intricately tied to ambient temperatures (Garrabou *et al*. 2009; Pinsky *et al*. 2019). Elevated temperatures exponentially amplify metabolic demands, thereby escalating the likelihood of bioenergetic failure (Pörtner *et al*. 2010). When temperatures exceed a critical threshold, a cascade of physiological disruptions ensue, affecting mechanisms from oxygen balance and cellular redox states to enzyme activities and membrane stability (Boutilier & St-Pierre 2000; Somero 2010; Pörtner 2012; Ern *et al*. 2023; Sokolova 2023). While these disruptions harm all life stages, they particularly jeopardize early life stages, such as larvae and juveniles, which are not only less physiologically resilient but also crucial for population renewal (Collin *et al*. 2021).

The resulting heightened mortality rates during heatwaves, especially among foundational species, can set off ecological cascades, threatening ecosystem stability. This is evidenced by the heatwave-induced die-off of vital ecosystems like coral reefs, seagrass beds, kelp forests, and mussel banks (Wernberg *et al*. 2016; Hughes *et al*. 2017; Seuront *et al*. 2019; Strydom *et al*. 2020). Such events can lead to broad shifts in species distributions and even biome restructuring (Bellard *et al*. 2012).

However, marine heatwaves also act as selective forces, favoring heat-tolerant individuals and offering a mechanism for evolutionary rescue (Gomulkiewicz & Holt 1995; Gilbert & Miles 2017; Vajedsamiei *et al*. 2021c). To address the pressing need for understanding the eco-evolutionary implications of marine heatwaves, computational models considering intra-species variability in heat sensitivity such as Thermal Tolerance Landscape TTL—as termed by (Rezende *et al*. 2014)—have emerged as indispensable predictive tools. These models quantify population survival based on temperature and duration of exposure (see Supplementary Text 1), positing that beyond critical temperature thresholds, the *lethality buildup rate* escalates exponentially but remains constant in time (Jacobs 1919; Bigelow 1921; Fry *et al*. 1946; Jørgensen *et al*. 2019, 2021; Rezende *et al*. 2020). Rationale for using the term lethality buildup rate is explained in Supplementary Text 2.

In this context, TTL models have been mainly parametrized using frequentist linear regression of survival times or the reciprocal—lethality buildup rates—over constant temperatures (Jørgensen *et al*. 2019, 2021; Rezende *et al*. 2020). This method can derive key parameters including the mean lethality buildup rate at the reference temperature (intercept) and thermal sensitivity parameter (slope), which in turn inform prediction of mean population survival trajectories during dynamic heatwaves (Rezende *et al*. 2020; Bertolini & Pastres 2021; Jørgensen *et al*. 2021; Li *et al*. 2023). Despite its merits, the frequentist framework falters when it comes to offering probabilistic predictions (Ellison 2004). Moreover, the premise of physiological consistency over time, even if validated in controlled laboratory environments, might not stand firm as organisms could exhibit varied heat sensitivities when exposed to gradual temperature increases leading up to more potent lethal temperatures (Helmuth *et al*. 2004).

This paper refines the predictive capability of TTL models in capturing real-world dynamics through the novel application of Approximate Bayesian Computation (ABC) for TTL parameterization, specifically tailored for data obtained during natural dynamic heatwave events. As a study organism, we selected the Baltic *Mytilus* mussels, pivotal in their ecological role. Our experimental protocol involved monitoring survival rates of juvenile and adult mussels under two distinct conditions: microcosm-simulated constant heatwave (CHW) and mesocosm-simulated near-natural dynamic heatwave (DHW) regimes. Post-heatwave, recruitment counts within the mesocosms were undertaken. Our methodological objectives encompassed:

i. **Model parameterization:** We adopt an ABC approach to determine posterior parameter distributions for TTL models and to generate posterior predictions for trajectories of individual lethality buildup and population survival for mussels. Analyzing TTL outcomes from both CHW and DHW experiments provides insights into the applicability of the approach and tests whether assumptions of physiological homogeneity and temporally constant lethality buildup rates are sufficiently robust to generalize across different environmental settings.
ii. **Projections:** Applying temperature projections for high CO_2_ emission scenarios, we provide projected lethality buildups in mussels during the century’s five warmest summer regimes, taking daily temperature variability into account. We also illustrate how these predictions can serve as indicators of potential declines in mussel recruitment rates, using the established correlation between observed recruitment and modeled lethality from the DHW experiment.

Our methodologies promise to enhance our predictive capacity for key marine populations in the face of intensifying heatwaves, addressing the urgent need for accurate modeling tools to develop early warning systems and inform ecosystem management and policy decisions (Pecl *et al*. 2017; Skirving *et al*. 2019), ultimately aiding in the conservation of ecosystems in our rapidly changing climate (Harley *et al*. 2006; Poloczanska *et al*. 2013).

## METHODS

### Data collection

#### The study system

We utilized the blue mussel from the *Mytilus edulis* complex in the Western Baltic Sea as our model organism due to its extensive reef formations and significant ecological and economic roles (Zippay & Helmuth 2012; Burge *et al*. 2016; Larsson *et al*. 2017). The Baltic Sea, experiencing rapid warming trends, provided an opportune setting for our predictive heatwave response models, supported by available future temperature projections (Reusch *et al*. 2018; Meier *et al*. 2021).

#### Constant heatwave (CHW) experiment

The CHW experiment evaluated mussel survival under constant heatwave conditions, targeting our first research objective. Mussels were exposed to temperatures of 26, 27, 28, or 29 °C, levels known to cause mortality within days to weeks (Zittier *et al*. 2015; Vajedsamiei *et al*. 2021c).

On July 6^th^, 320 mussels were collected from a location near GEOMAR at less than 0.2 meters depth (54°19’45.4″N 10°08’56.2″E), sorted into two size classes (approximately 7 and 20 mm), and acclimated for a week in 8-L aquaria at 17°C. They were fed daily with *Rhodomonas salina* (initial concentration of 1875 cells mL^-1^) cultured at GEOMAR’s KIMOCC, with water changes to mitigate ammonia.

From July 12-15, we set up eight thermal baths using the Kiel Indoor Benthocosm (KIB) system, featuring computer-controlled 600-L tanks (Pansch *et al*. 2018). In each, four 18-L cylinders held 1 μm filtered seawater: one unaltered, one with a temperature probe, and two containing 2-L PVC baskets (5-mm mesh).

On July 15, within a 10-minute window starting at 12:00 am, 320 mussels were evenly distributed across 16 PVC baskets (80 per temperature, 40 per bath). Temperature was recorded hourly by the GHL system (Profilux 3.1TeX; GHL Advanced Technology) and verified daily with a handheld sensor (WTW Multi 3630 IDS), which also measured water salinity, pH, and oxygen on July 15 and weekly thereafter. Mussels were moved to fresh, temperature-equilibrated seawater every five days, and water in used cylinders was replaced.

Mortality was assessed daily at a consistent time, using unresponsiveness (no valve gaping) under mechanical stress. Dead mussels were stored in plastic bags until the experiment concluded on August 25, at which point their shell lengths were measured. Additional details on the setup and measurements can be found in Supplementary Figures 1 and 2.

#### Dynamic heatwave (DHW) experiment

The DHW experiment was conducted to evaluate mussel survival and recruitment under dynamic heatwave conditions.

480 mussels collected on July 1 were maintained in two 17°C aquaria for a week, following the same protocol as the CHW experiment. On July 6, these mussels were equally distributed across 12 PVC cages, each containing a mix of 20 small and 20 large sized mussels. These cages were then installed in the twelve 1400-L tanks of the Kiel Outdoor Benthocosm system (KOBs; referenced in Wahl *et al*. 2015). Three peristaltic pumps distribute non-filtered seawater (40,000 L d^-1^) from the fjord to sets of four tanks. The computer-controlled units mimic near-natural temperature dynamics in the tanks where water is fully mixed due to built-in circulation and a wave generator.

Tanks in the experiment had dynamic temperature baselines, staggered from 0°C to 5.5°C at 0.5°C intervals (see Supplementary Figure 1). From July 7 to 13, temperatures in eleven tanks were adjusted to establish these baselines. We simulated a heatwave from July 29 to September 1 by reducing inflow to 4000 L d^-1^ and increasing baseline temperatures by 2.5°C. Adjustments occurred between 9:00 and 10:00 am, with the baseline returning to ambient Kiel Fjord temperatures on September 1. Temperature, O_2_, pH, and salinity were recorded daily, and chlorophyll levels were measured weekly, as detailed in Supplementary Figures 3, 4, and 5.

By the end of the DHW experiment (September 8), mussel survival and sizes were assessed using the methods described for the CHW experiment. To evaluate mussel recruitment, settlement panels were installed on June 2 in all 12 tanks near but not in direct disposal of the wave generator and above the *Fucus* habitats so that larval mussels could attach, metamorphose and grow in the absence of predation. These panels consisted of four types: Two PVC (0.3×6×6 cm and 0.5×6×6 cm) and two brick (3×3×3 cm and 3×6×6 cm). Recruitment data were collected at the end of the experiment, with settled mussels (1 to 10 mm final shell length) collected and counted.

#### Data processing and modelling

The Thermal Tolerance Landscape (TTL) equations are outlined in Supplementary Text 1. Briefly, Equation 4 quantifies the cumulative lethal heat stress that an organism experiences (*lethality buildup*; see Supplementary Text 2) when exposed to consistent or varying temperatures over time. It does this by integrating the lethality buildup rates, while each rate being adjusted depending on the difference between the reference and actual temperature experienced by the organism (Equation 7). The integrated lethality can then be used to estimate population survival under various temperature conditions, as outlined by Equations 1–3.

The subsequent section details an Approximate Bayesian Computation (ABC) to generate the posterior distributions of three TTL parameters: the temperature sensitivity parameter, and the mean and standard deviation of the lethality buildup rate at the reference temperature, i.e., respectively denoted as, k, mean 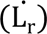, and 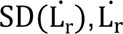 on the decadic logarithm scale. The reference temperature (T_r_) is set as 28°C. This approach utilizes data independently derived from the CHW and DHW experiments.

In Supplementary Text 4, an auxiliary Bayesian Regression (BR) approach is also described to analyze the lethality buildup rate data—essentially, the reciprocal of survival times—from the CHW experiment. Although BR is not the primary methodology of this paper, it represents an enhancement over the previously utilized frequentist regression for TTL parametrization. Thus, we also compare its outcomes with the results from the ABC methodology outlined below.

The computational methods used are written and executed in the *R* programming language (version 4.2.3), with all scripts being accessible on GitHub (https://anonymous.4open.science/r/A-Bayesian-overhaul-of-thermal-tolerance-landscape-models-17F7/). It is noteworthy that in both CHW and DHW experiments, survival patterns did not significantly vary between mussels sized 7 and 20 mm (refer to Supplementary Figure 6 and Supplementary Table 1). Consequently, the combined data set was utilized for further analysis.

#### Approximate Bayesian Computation (ABC)

We present an approach utilizing ABC with Sequential Monte Carlo (ABC-SMC; Sisson *et al*. 2007; Del Moral *et al*. 2012) for estimating posterior distributions of TTL parameters.

ABC is a specialized form of Bayesian analysis employed when calculating the likelihood—that is, the probability of observed data given certain parameters—is difficult or impossible while data simulation is feasible (Supplementary Text 3 provides a historical introduction to ABC). The ABC-SMC approach is based on a multi-iteration algorithm to refine parameter estimates progressively.

Figure 1 illustrates a schematic overview of our ABC-SMC methodology. The initial step involves sampling 5000 particles, with each representing a potential set of the three model parameters, from Gaussian prior distributions. In this study, while we inform these priors using posterior parameter distributions derived from the BR analysis on the CHW experiment data (as elaborated in Supplementary Text 4), this approach is not mandatory. The priors are designed with wider standard deviations to encompass a broader parameter space. It is worth noting that the ABC-SMC algorithm does not necessitate BR-based priors; preliminary runs of the algorithm itself can suffice for generating initial parameter estimates. These preliminary estimates are then iteratively refined through the ABC-SMC process, which can converge to yield robust results that are comparable to those obtained when Bayesian-informed priors are used.

**Figure 1.**
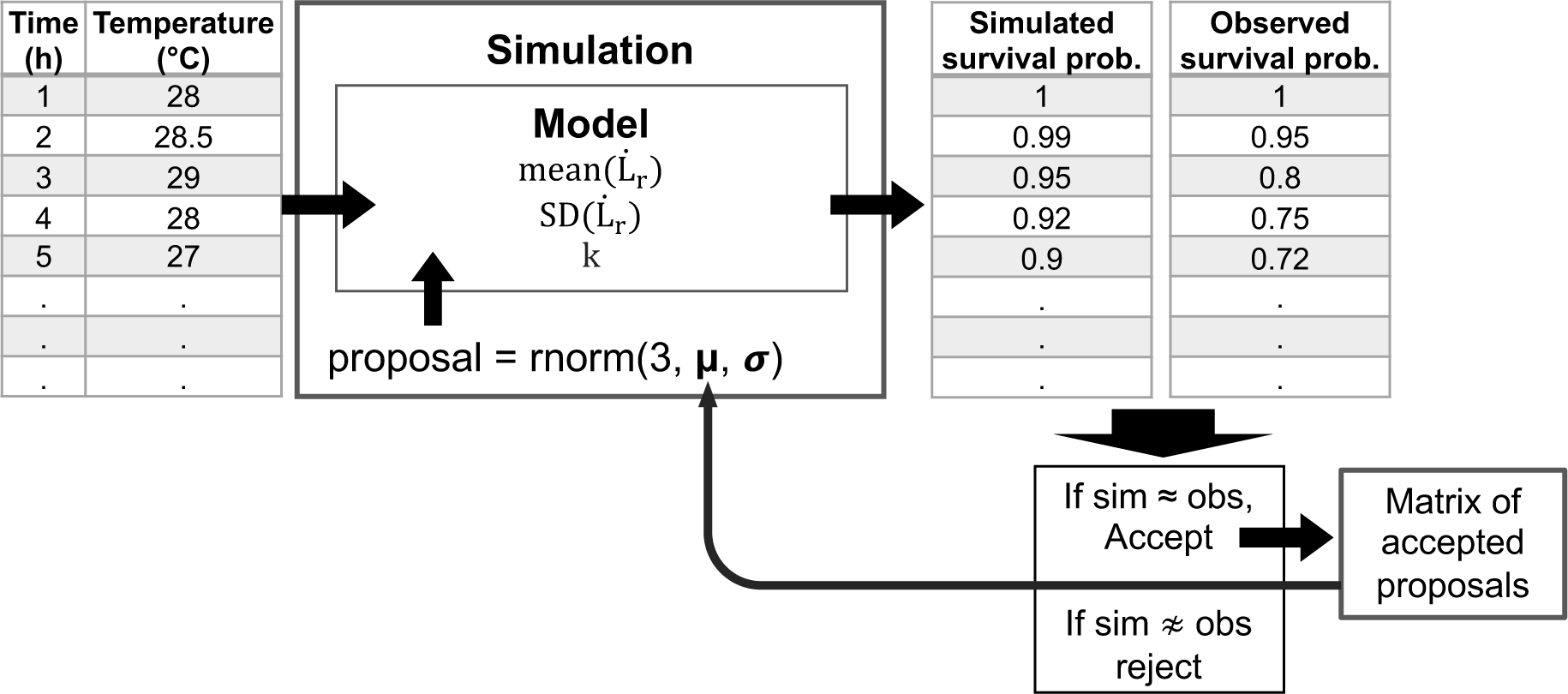
Schematic representation of the Approximate Bayesian Computation via Sequential Monte Carlo (ABC-SMC) for refining parameters of the Thermal Tolerance Landscape (TTL) model. The process begins with temperature time series data input into the TTL model alongside parameter proposals randomly drawn from a normal prior distribution, indicative of prior beliefs about the parameters’ likely values. The model iteratively simulates survival probabilities, which are compared to observed data using the Mean Absolute Deviation (MAD) across all temperature series within an experiment for each proposal. Proposals yielding simulated probabilities with MAD within an acceptable range are retained, contributing to a matrix that incrementally refines the posterior parameter distributions. With each iteration, the acceptance threshold narrows, integrating accepted parameters to calibrate subsequent simulations and enhance estimate precision. For an in-depth explanation of the methodology and the script, refer to Supplementary Text 5. Displayed values are illustrative.

In the initial phase of our algorithm, each particle is input into the TTL model to simulate population survival trajectories. We retain a particle if its Mean Absolute Deviation (MAD) from the observed survival data across all four CHW or DHW samples remains under the set initial MAD threshold (epsilon). These qualifying particles are then given equal probability for resampling in the subsequent iteration. Our approach iteratively refines the particle set through cycles where particles are resampled based on their weights, perturbed by adding Gaussian noise to them, and only retained if they fulfill the MAD criterion. If a particle fails to produce valid data after ten attempts, it is discarded. The algorithmic process concludes when the particle acceptance rate drops below 20%, resulting in a particle set that approximates the posterior distribution. This set offers insights into the most probable parameter values given the observed data. For an extensive discussion of the algorithm, refer to Supplementary Text 5.

To assess the performance of the ABC-SMC approach, we contrast the posterior predictions of mussel survival rates with the observed survival rates visually and estimate the MADs across various treatment levels of both CHW or DHW experiment. We compare the approximate posterior distributions between the two experiments using density graphs to evaluate the effects of varying environmental conditions. Supplementary Text 6 details the processes for posterior prediction (simulations) and the comparisons of MADs between the ABC-SMC and BR approaches.

Additionally, we draw the most likely posterior parameter set (defined by ABC-SMC) and visualize the most likely lethality buildup trajectories and the population survival trajectory under various regimes of DHW experiment, with an emphasis on delineating sublethal effects induced by heatwaves.

#### Projections of lethality buildups and survival rates in mussels

To prepare the data for these analyses, we adopted bi-daily sea surface temperature projections for the 21^st^-century Baltic Sea under the Representative Concentration Pathway 8.5 scenario, as projected by Meier et al. (2021). We refined these data to an hourly scale through linear interpolation, focusing on the five warmest summers near the Kiel Fjord entrance (Coordinates: 54°26’55.9″N 10°15’45.2″E). Additionally, we constructed a daily temperature fluctuation cycle on an hourly scale. This was informed by data collected from a shallow-water habitat in Kiel Fjord (1m depth; Coordinates: 54°23’44.6″N 10°11’27.5″E) during summer 2019. The mean daily fluctuation was calculated by averaging the differences between the actual and mean daily temperatures. These fluctuation patterns were then integrated into the temperature projections (see Supplementary Figure 7).

Inputting the temperature series of each summer (both inclusive and exclusive of daily fluctuations) to the TTL model, optimized using ABC-SMC on DHW data, we simulated both lethality buildup rates and corresponding population survival trajectories.

#### Average ABC-predicted lethality buildup as indicator of recruitment rate

Acknowledging the difficulties of tracking early-life-stage individuals in mesocosm experiments, we adopt the average (ABC-predicted) lethality buildup at the summer end as a representative indicator of the recruitment decline.

We calculate the average buildup for each mussel group in the DHW tanks and used a Bayesian regression model with the *brms* package (Bürkner 2017). The model, featuring a spline function with a Gamma-distributed error and a log-link response, ran on four chains of 10,000 iterations, including a 5,000-iteration warmup. With the model constructed, we generate normalized recruit abundance predictions over a spectrum of average lethality buildups, forming the basis for our recruitment projections.

## RESULTS

In constant heatwave (CHW) regimes, survival of mussels (initial length range of ca. 7-20 mm) reached zero within 7 days at 29°C and within 30 days at 28°C. At lower temperatures, survival probabilities were approximately 0.24 at 27°C and 0.7 at 26°C at the end of the 45-day experimental period. In the dynamic heatwave (DHW) experiment, a notable decline in mussel survival was evident exclusively in the four most elevated temperature regimes. Survival rates declined to zero within a period of 52 days at the highest temperature regime, which was characterized by a median temperature of 25.2°C and a minimum-maximum range of 19.1-28.8°C. At the end of the 64-day experimental period, survival rates were documented at 0.875, 0.825, and 0.325 at median temperatures of 23.8°C (with a min-max range of 19.4-27.4°C), 24.3°C (19.6-28.1°C), and 24.8°C (20.3-28.6°C), respectively. All twelve temperature regimes employed in the DHW experiment are presented with varied temperature statistics, as detailed in Supplementary Table 2.

### Robust parameter estimation and validation of TTL models using ABC-SMC

Utilizing Approximate Bayesian Computation via Sequential Monte Carlo (ABC-SMC), we achieved precise parameter estimations for Thermal Tolerance Landscape (TTL) models. The model-generated survival probabilities closely aligned with the 95% confidence intervals of Kaplan-Meier curves from both CHW and DHW experiments (Figure 2). In assessing model fidelity, we have achieved a noteworthy level of accuracy through the assessment of Mean Absolute Deviance (MAD) values, which quantify the disparity between survival predictions simulated by the ABC-SMC-parametrized model and the corresponding observed data. Across all simulated populations, the median MAD for the CHW condition was calculated to be 0.0572, with a 5^th^-95^th^ percentile range of 0.0340 to 0.0758. Similarly, for the DHW condition, the median MAD was found to be 0.0203, with a 5^th^-95^th^ percentile range of 0.0104 to 0.0371. These remarkably low MAD values, coupled with the narrow range of variability, provide strong evidence of the accuracy and reliability of the ABC-SMC model.

**Figure 2.**
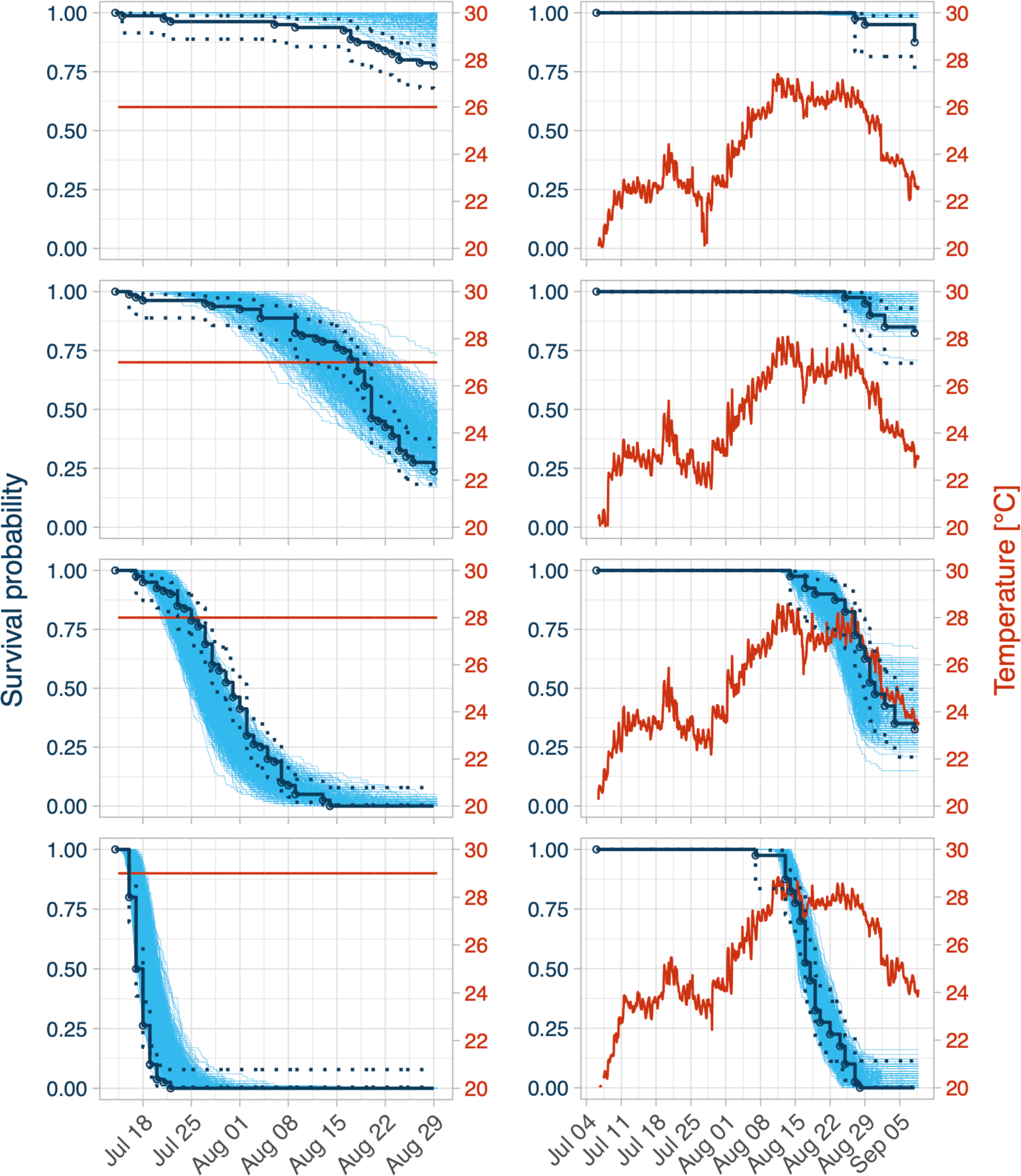
Mussel survival rates at constant and dynamic temperature regimes (left versus right-side columns). Trajectories simulated based on posterior parameter sets from the ABC-SMC are presented as bright blue lines. The observed survival probabilities are presented as black circles with the corresponding interpolated Kaplan-Meier predictions (continuous black lines) with 95% confidence interval (dotted lines). Thermal regimes are displayed in red.

Additional insights into the performance of the ABC-SMC model in comparison to the Bayesian Regression (BR) approach are presented in Supplementary Figure 10 and Supplementary Table 3. Overall, both methods demonstrated comparable performance in the context of the CHW experiment, yielding consistently low MAD values. However, in the case of the DHW experiment, as anticipated, the ABC-SMC approach outperformed BR by achieving higher accuracy. This outcome is expected, given that the ABC-SMC model was parameterized directly using data collected under DHW conditions. In contrast, the BR approach yielded a median MAD of 0.0469 for DHW, with a 5^th^-95^th^ percentile range of 0.0370 to 0.0577, which is notably larger than the MAD obtained through ABC-SMC.

Figure 3 showcases the approximate posterior distributions of the TTL parameters. The bell-shaped (Gaussian) distributions affirm successful sampling of the parameter spaces. Notably, the overlap within the 95% credible intervals (CI) of the pair distributions suggests that differences in the TTL parameters in response to varying temperature regimes were not statistically significant (Figure 3). Yet, the difference in the mean value of the parameter *lethality buildup rate standard deviation* 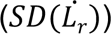 between the CHW and DHW conditions was relatively more pronounced, especially on the log_10_-transformed scale due to its compressive effect (Figure 3; Supplementary Table 4). This manifested as the SD of the log_10_-transformed 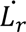 appearing 33% higher under constant conditions, a distinction that diminishes when reverted to the original scale, where the constant regime’s SD was approximately 19.8% higher. On the original scale, there was, on median, a 6% increase in the mean of 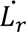 from the constant to the dynamic experiment, suggesting a slightly faster lethality buildup at 28°C under the dynamic regime. The parameter k, denoting temperature sensitivity of lethality buildup rate, also increased by roughly 13% under dynamic conditions, suggesting higher responsiveness to temperature changes.

**Figure 3.**
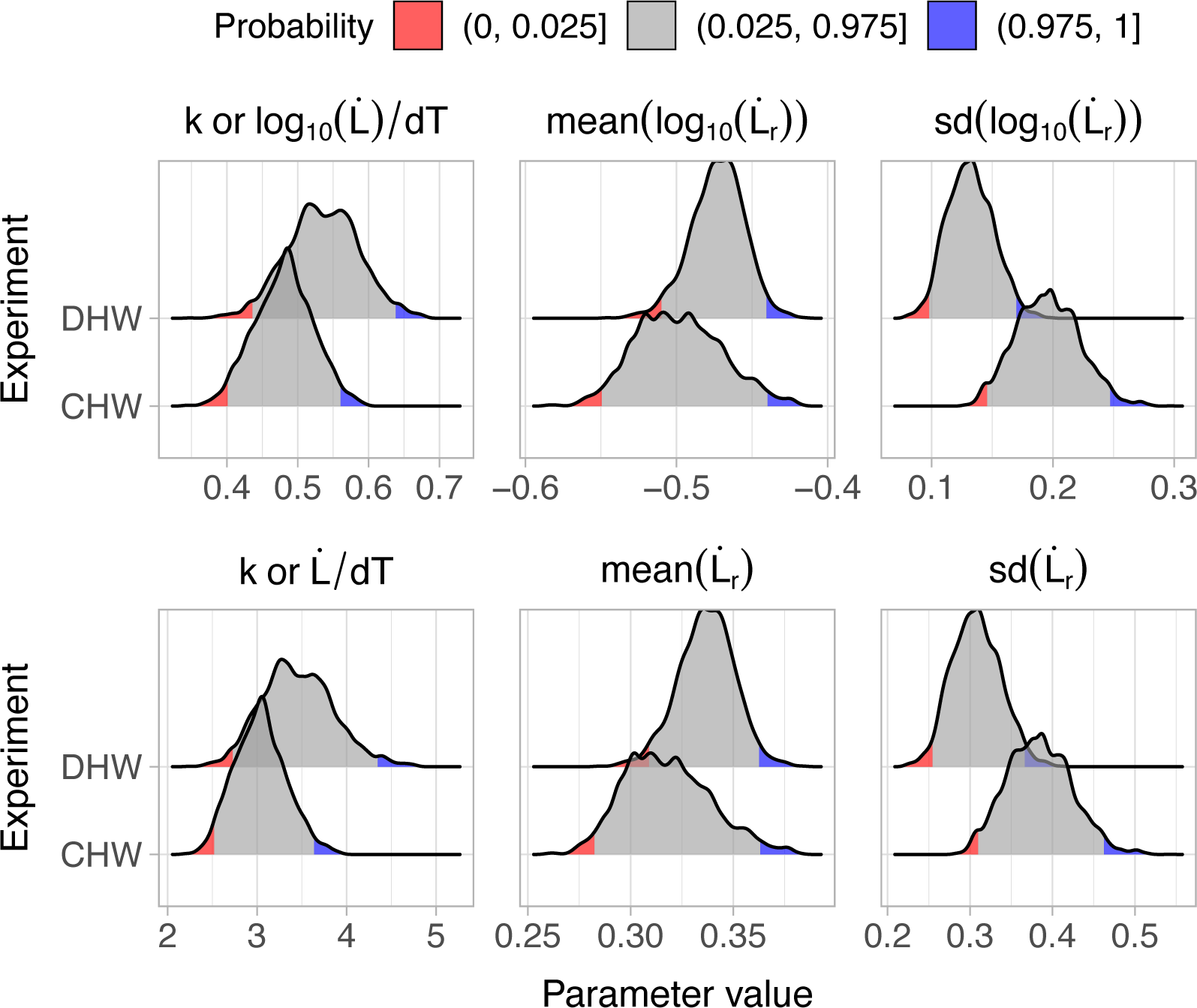
Approximate parameter distributions for the TTL model derived using Approximate Bayesian Computation via Sequential Monte Carlo (ABC-SMC), informed by experimental data from constant (CHW) and dynamic (DHW) heatwave experiments. The subplots, arranged from left to right, illustrate the distributions of the temperature sensitivity parameter *k*, and the mean and standard deviation of the lethality buildup rate at the reference temperature [*mean*(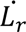) and *SD*(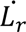), respectively], with 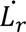 expressed on the decadic logarithm scale. The 95% credible interval is represented in grey, and the lower and upper 2.5% probability quantiles are shown in red and blue, respectively.

Figure 4 demonstrates the trajectory of population survival and lethality buildups for individual mussels under all dynamic temperature regimes. These trajectories, demonstrated as light blue and green lines respectively, were generated based on the most probable posterior parameter set derived from the ABC-SMC approach. The applied temperature regimes, indicated by the red lines, embody a wide range of thermal conditions under which we assessed mussel survival. Based on the TTL, the survival probability of the mussel population is predicted to decrease in response to the three warmest treatment conditions. However, sub-lethal effects in mussel individuals (i.e. scaled lethality buildup<1) is predicted to occur across more temperature treatments, not just the warmest ones. Over the 5 warmest DHW regimes, nearly all individuals of the populations were predicted to buildup > 0.1 or >10 % lethality.

**Figure 4.**
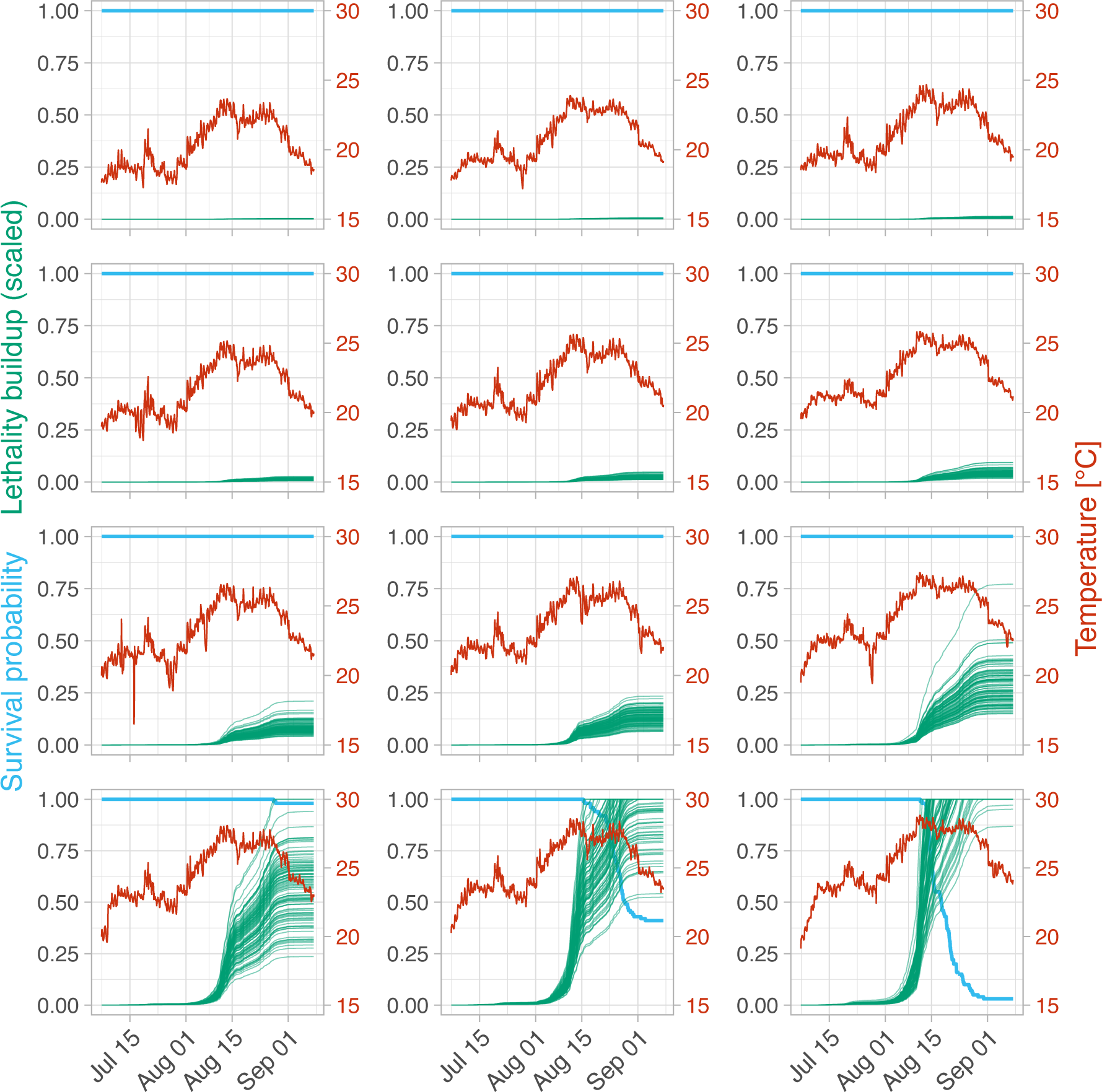
Simulated survival (light blue lines) and scaled lethality buildup (green lines) trajectories for a mussel population subjected to twelve dynamic heatwave (DHW) regimes (shown in red; temperature regimes’ statistics presented in Supplementary Table 2). The simulations are based on the most likely posterior parameter set, obtained via Approximate Bayesian Computation (see the main text), using observed survival rates from the DHW experiment. The corresponding thermal regimes are depicted in red.

### Projections of mussel lethality and survival rates

Our projection analysis, utilizing the TTL with most likely parameter set obtained using ABC on data from the DHW experiment, indicates that mussels (with shell lengths of 7-20 mm in early July) in the western Baltic Sea’s subtidal habitats are not expected to experience mortality under any of the five warmest summer scenarios projected for this century. This is consistent across all scenarios, regardless of the inclusion of daily fluctuations (as shown by the blue lines in Figure 5). However, each scenario does contribute to an increasing lethality within the mussels (illustrated by the green lines in Figure 5; see also Table 1). Notably, the comparison of scenarios with and without daily fluctuations (right versus left panels in Figure 6) underscores that daily temperature fluctuations may potentially amplify the harmful impacts of warmest summer heatwaves on lethality buildup in young mussels.

**Figure 5.**
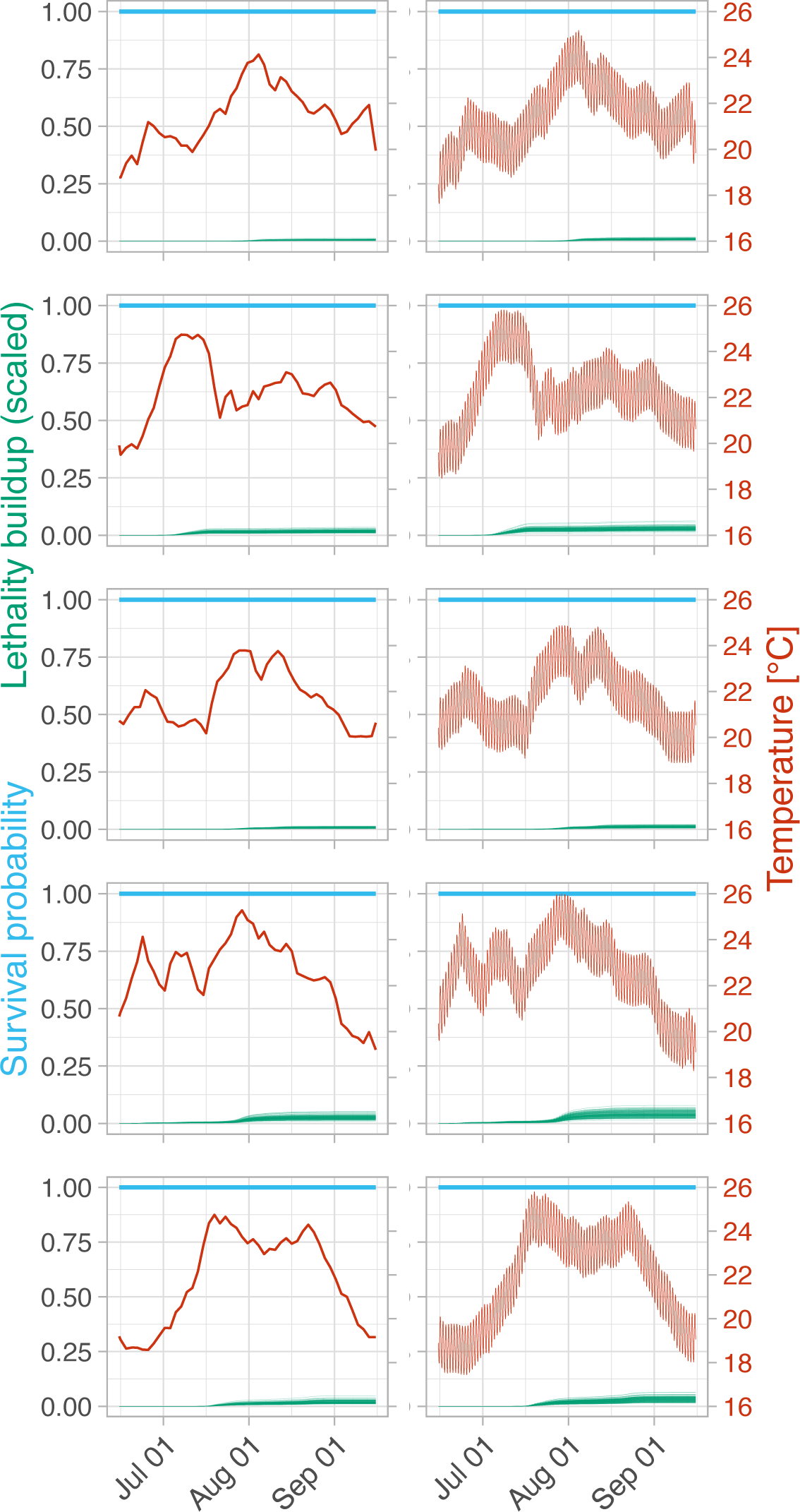
Projected survival and mean lethality buildup for mussels under five warmest summer temperature projections for the years 2060, 2083, 2085, 2092, and 2094 (top to bottom), considering scenarios with and without daily temperature fluctuations (left versus right-side subplots).

**Table 1.** Projected average lethality buildup (Average L) and recruitment probabilities (with the 50^th^, 95^th^, and 5^th^ percentiles) under the five warmest projected summer regimes, with and without daily fluctuations.

| Year | Fluctuation | Average<br>L | Scaled<br>recruitment<br>P50 | Scaled<br>recruitment<br>P5 | Scaled<br>recruitment<br>P95 |
| --- | --- | --- | --- | --- | --- |
| 2060 | yes | 0.009 | 0.989 | 0.647 | 1.471 |
|  | no | 0.006 | 1.000 | 0.658 | 1.497 |
| 2083 | yes | 0.028 | 0.775 | 0.497 | 1.144 |
|  | no | 0.018 | 0.893 | 0.583 | 1.326 |
| 2085 | yes | 0.011 | 0.973 | 0.636 | 1.444 |
|  | no | 0.007 | 0.995 | 0.658 | 1.487 |
| 2092 | yes | 0.04 | 0.658 | 0.422 | 0.968 |
|  | no | 0.026 | 0.797 | 0.519 | 1.182 |
| 2094 | yes | 0.032 | 0.733 | 0.471 | 1.086 |
|  | no | 0.022 | 0.840 | 0.545 | 1.246 |

### Logistic decline in mussel recruitment under elevated DHW regimes

The DHW experiment showed a logistic decline in the abundance of recruited (settled) mussels with the warming, with a sharp decrease at medium DHW levels (3 °C above baseline), leading to near-zero recruitment at the highest DHW (see Supplementary Figure 11, left side plot). This pattern is supported by our Bayesian regression model’s precise fit to the data. Supplementary Figure 11 displays this association and, alongside, shows the correlation between observed recruitment and predicted lethality buildup in older mussels (ABC-L in both logarithmic and original scales) subjected to the same DHW regimes.

The latter correlation—though not indicative of a causal relationship—suggests that the low values of average lethality buildup projected for the five most extreme future summer regimes may signal potential declines in recruitment probability (Table 1). Projections indicate more declines in median recruitment under scenarios with daily fluctuations compared to those without (as detailed in Table 1), while the large credible intervals point to significant variability in recruitment rates within our KOB system (Supplementary Figure 11, left side plot). For instance, in the year 2094, we project the average lethality buildup in juvenile mussels to be 0.032 under fluctuating conditions and 0.022 without fluctuations. These projections translate to median recruitment probabilities of 0.733 (95% credible interval: 0.471–1.086) and 0.840 (95% CI: 0.545– 1.246), respectively. Notably, these projections assume that the median recruitment rate (Supplementary Figure 11, middle plot) is capped at an optimum value of one and that the credible intervals are scaled proportionally.

## DISCUSSION

Our study has refined Thermal Tolerance Landscape (TTL) modeling by an innovative application of Approximate Bayesian Computation-Sequential Monte Carlo (ABC-SMC) to analyze survival in Baltic *Mytilus* mussels under heatwaves. This approach yields probabilistic assessments of sublethal impacts and mortality, promising a broad application for ectotherms confronting climate change.

### TTL parametrization via ABC: Key for predictive insights on heatwave-induced lethality in ectotherm populations

ABC-SMC has enabled precise TTL parameter estimation from mussel survival data under constant and dynamic heatwave (CHW and DHW) regimes, validated by low MAD values when compared to observed rates. This substantiates the TTL assumption that survival times logarithmically decline with increasing temperatures, adhering to foundational thermal tolerance research (Jacobs 1919; Bigelow 1921; Rezende *et al*. 2020), and confirming that lethality buildup rates rise exponentially with temperature while remaining stable at constant temperatures (Fry *et al*. 1946; Jørgensen *et al*. 2019, 2021), with minimal compensatory acclimation (Havird *et al*. 2020; Vajedsamiei *et al*. 2021b).

Prior to our study, TTL models were primarily parametrized using frequentist linear regressions, relying on constant temperature experiments (Rezende *et al*. 2020; Jørgensen *et al*. 2019, 2021; Bertolini & Pastres 2021; Li *et al*. 2023). Our study advances this by applying a Bayesian regression (BR) within the TTL framework, which not only predicts average survival trajectories but also provides a probabilistic understanding of parameter distributions. Detailed in Supplementary Text 4, the BR approach streamlines the Monte Carlo simulations of potential survival trajectories.

Robust TTL modeling necessitates incorporating data from a spectrum of temperatures and must consider censored data where not all individuals die within the experimental timeframe. Our results showed that both the BR and ABC-SMC techniques accurately estimate survival rates under CHW conditions, accounting for data censoring.

Predicting responses to DHW regimes using a TTL model parameterized by CHW experiment data is problematic. The variance in MAD values between BR and ABC-SMC methods demonstrates the complexity of DHW scenarios, as gradual exposures to temperatures above 24°C can affect mussels differently compared to CHW exposures (Vajedsamiei *et al*. 2021a; Khosravi *et al*. 2023). More accurate prediction of population survival under DHW regimes demands the application of ABC-SMC on data derived from DHW experiments.

Our approach additionally enabled the prediction of probable trajectories of accumulating sublethal effects during heatwaves on mussels. Ectotherms like *Mytilus* mussels exhibit intricate, temperature-sensitive physiological responses, which encompass modifications in metabolic rates, enzyme activities, cellular homeostasis, and gene expression patterns (Connor & Gracey 2020; Georgoulis *et al*. 2022). Under heightened temperatures, these alterations can lead to a range of physiological setbacks, such as weakened immune functions (Cellura *et al*. 2007; Hong *et al*. 2021), reduced growth rates (Vajedsamiei *et al*. 2021b) and perturbed reproduction (Béjaoui-Omri *et al*. 2014). The sublethal lethality buildup modeled in our study can quantitatively represent these responses to stress and fitness setbacks.

### Projections of lethality buildup and survival in mussels

Our TTL projections—derived from the most probable parameters obtained through ABC-SMC applied to DHW experiment data—indicate that juvenile and adult mussels (shell length 7-20 mm) in the western Baltic Sea’s subtidal zones may demonstrate resilience against the most extreme summer temperatures expected this century. However, we predicted some sublethal effects, exacerbated by substantial daily temperature fluctuations, which underscores the need for attention regarding potential carryover effects in the next seasons and echoing prior evidence on the importance of diurnal temperature variability in marine species’ heat resilience (Vasseur *et al*. 2014; Vajedsamiei *et al*. 2021b).

For clarity in our methods, we note that our method practice primarily projected the repercussions expected in the five warmest summers by 2100, specifically under the RCP 8.5 climate scenario, and considered one extensive daily fluctuation level, occurring in shallow waters (less than 1 meter deep) with gradual inclines. Yet, our data, predominantly from open and deeper subtidal areas, may not fully capture the resilience and thermal conditions of mussels in more restricted shallow waters, where higher peak summer temperatures prevail. Acknowledging this data gap in key habitats, it becomes imperative to broaden our research scope and heat tolerance phenotyping.

### Logistic decline in recruitment under elevated DHW regimes

Our study has revealed a more pronounced heat sensitivity of *Mytilus* during the recruitment phase, compared to older juveniles and adults. Considering the developmental period of Baltic *Mytilus* larvae—typically lasting 3-5 weeks post-peak spring spawning—and the rapid water exchange rate in our mesocosm tanks, the settled larvae were evidently sourced externally (Seed 1969; Thorarinsdóttir & Gunnarsson 2003; Büttger *et al*. 2011). Therefore, the observed decline in recruit abundance can be attributed to the adverse thermal effects on incoming larvae from the fjord during their critical final metamorphosis phase (late pelagic larvae) or on newly settled juveniles. Further detailed research is essential to identify the specific stages and processes that contribute to the high thermal sensitivity observed in mussel recruitment.

The literature corroborates that temperature extremes impact early life stages of ectotherms, with implications for the population dynamics and long-term survival of marine species under increasing heatwave conditions (Sorte *et al*. 2011; Collin *et al*. 2021). Nonetheless, tracking survival rates of settling mussels over time—needed for TTL modeling and predictions—presents a significant challenge in mesocosm studies. However, using a parametrized TTL model to quantitatively predict lethality buildup in juvenile and adult mussels offers an indirect but insightful way to project recruitment trends. This is particularly useful since lethality buildup acts as an indicator for systemic stress, and both recruitment and lethality are driven by temperature. Building on this correlation, our projections indicate that under the century’s top-five most extreme temperature scenarios expected by 2060, recruitment could decline by as much as 0-20% in the absence of daily temperature fluctuations, and between 1-27% when considering such fluctuations. These findings highlight the potential severity of heatwaves on mussel populations.

With the ongoing shift in global temperatures, mussels may undergo eco-evolutionary changes, including phenological shifts that could alter recruitment timing (Filgueira *et al*. 2015). While not included in our current model, such changes are critical to the future dynamics of mussel populations.

## Conclusion

Our study applied the ABC-SMC method to enhance TTL modeling for Baltic *Mytilus* mussels, demonstrating superior accuracy and applicability over traditional frequentist and Bayesian regression methods. The ABC-SMC method effectively identified the most likely parameter sets from survival data under various heatwave conditions. The capacity of ABC-SMC to utilize data from mesocosm studies, which closely mirror natural habitats, proved invaluable. Our results also indicated that TTL model-projected lethality buildup could serve as an early warning signal for potential declines in recruitment amidst intensifying heatwaves.

Envisioning the progression of this field, we propose several research directions:

1. Delineating the precise relationships between physiological responses and lethality buildup amidst heatwaves remains an area for future research, essential for decoding the temporal dynamics of mechanisms that culminate in mortality during such events.
2. Extending these methods to foundational coastal species, thereby equipping us with predictive tools to measure how key marine populations might respond to escalating heatwaves. This could play a crucial role in developing strategies to mitigate the impacts of climate change on essential marine ecosystems.

