## Supplementary Materials for "A Bayesian overhaul of thermal tolerance landscape models: Predicting ectotherm lethality buildup and survival amid heatwaves"

**Contents:**

**Supplementary Texts**

**Supplementary Figures**

**Supplementary Table**

### Supplementary Texts

#### Supplementary Text 1. Principles of Thermal Tolerance Landscape (TTL) model

The principles of the TTL model are outlined in the following equations, defining how population survival rates change over time and temperature.

*Survival probability equations:*

Eq. 1. $\mathrm{P(t)} = 1 - \frac{\mathrm{M_p(t)}}{\mathrm{n_p}}$

Eq. 2. $\mathrm{M_p(t)} = \sum_0^t \mathrm{M_i}$

Eq. 3. $\mathrm{M_i} = \{\, 1, \text{if}\, L_i \;=\; 100;\; 0, \text{otherwise} \,\}$

Here, $P(t)$ represents the population's survival probability at time t. $M_p$ indicates the total mortality within the population up to time t, and $n_p$ is the population size. Individual mortality, denoted by $M_i$, is set to 1 when the accumulated lethality, $L_i$, reaches 100.

*Lethality buildup equations:*

Eq. 4: $L_i(\mathrm{t}) = L_{i_0} + \int_0^{\mathrm{t}} \dot{L_\iota}(\mathrm{t})\,\mathrm{dt}$

In this equation, $L_i(t)$ signifies the lethality buildup for an individual at time t, with $\dot{L_\iota}(\mathrm{t})$ denoting the rate at which lethality builds up at that specific moment. It is also assumes there is no initial lethality buildup present prior to the heat wave exposure, $L_{i_0}$= 0.

*Rate of lethality buildup equations:*

Eq. 5: $\dot{L_\iota} = \frac{1}{\mathrm{t}_i}$

Eq. 6: $\dot{L_\iota} = 10^{k_p(\mathrm{T-c})}$

Eq. 7: $\dot{L_\iota}(\mathrm{t}) = L_{i_r} \times 10^{k_p(\mathrm{T(t) - T_r})}$

Equation 5 calculates an individual's lethality buildup rate based on its survival time at a constant temperature. As an individual's survival time decreases, the lethality buildup rate accelerates. Equation 6 represents the relationship between lethality buildup rate and temperature ($T$), with parameter $k_p$ indicating the shared population-level sensitivity to temperature changes and constant c adjusting the temperature scale. With some algebraic manipulation, Equation 7 is reached that adjusts the lethality buildup rate based on the difference between a reference temperature ($T_r$; an arbitrary chosen lethal temperature) and the actual temperature over time $T(t)$.

#### Supplementary Text 2. Rationale for using the term *lethality buildup rate*

This study uses the term *lethality buildup rate* as a concise adaptation of the *lethal experience accumulation rate* from Fry *et al.* (1946). The choice of *buildup* rather than *accumulation* is deliberate; it underscores the escalating danger as temperatures increase—reflecting a linear increase of lethality over time and an exponential increase with temperature. This term effectively communicates the enhanced risk without presupposing specific physiological or physical mechanisms of death, which frequently remain unidentified. Thus, *lethality buildup* offers broader applicability, focusing on the end result rather than the cause. In contrast, terms like *damage accumulation rate* or *injury accumulation rate*, applied by Jacob (1929) and Jorgensen *et al.* (2021), are less neutral and not necessarily related to mortality.

**Supplementary Text 3. ABC: Origins, extensions, and applications**

Approximate Bayesian Computation (ABC) encompasses statistical techniques that leverage stochastic simulation for data-driven inference in complex models, especially with intractable or computationally demanding likelihood functions (Lopes & Beaumont 2010). Initially designed for population genetics, its applications have since burgeoned across fields like ecology, epidemiology, finance, and physics. Rubin (1984) was among the pioneers who proposed simulation-based Bayesian inference, which was subsequently advanced for parameter estimation and population structure inference (Tavaré *et al.* 1997; Pritchard *et al.* 1999). Innovations such as Markov Chain Monte Carlo (MCMC-ABC; Marjoram *et al.* 2003) sought to better approximate posterior distribution sampling, which was complemented by the regression adjustment (Beaumont *et al.* 2002). Sequential Monte Carlo (SMC) techniques, such as SMC-ABC (Sisson *et al.* 2007; Toni *et al.* 2008) and Adaptive SMC-ABC (Del Moral *et al.* 2012), improve sampling efficiency by employing intermediate distributions for more efficient parameter space exploration. Recent applications of ABC in biology demonstrate its versatility, with ABC being used to improve individual-based models (van der Vaart *et al.* 2016) and prey-predator models (Fasiolo & Wood 2018). Recently, deep ABC has emerged, utilizing deep learning for concise summary statistics learning and model selection (Lueckmann *et al.* 2017; Mondal *et al.* 2019). The versatility of ABC is demonstrated in a variety of biological applications, shedding light on numerous biological phenomena.

**Supplementary Text 4. Bayesian regression (BR) analysis on data from constant heatwave (CHW) experiment**

We provide an *R*-script for the BR analyses (refer to https://anonymous.4open.science/r/A-Bayesian-overhaul-of-thermal-tolerance-landscape-models-17F7/). Using Equation 5 (refer to the main text), we calculate the individuals' lethality buildup rates ($\dot{L}$) based on observed survival times at four constant temperatures (26, 27, 28, and 29°C). At 26 and 27 °C, the duration of the experiment was not long enough to cause 100% mortality. Therefore, the survival time data and $\dot{L}$ are 'censored,' meaning they are not recorded for a part of the mussel samples. Bayesian log-linear regression analysis of censored data is done using the *brm* function of the package *brms* (Bürkner 2017) to model $\dot{L}$ as a function of temperature. The model have an additional parameter, the censoring parameter (*cen1*), which represents the threshold below which the lethality buildup rate data are considered censored. For the fixed effects (intercept, slope, censoring parameters), the default priors are improper flat prior distributions. For the residual standard deviation, the default prior is a half *Student-t* distribution. The *brm* was applied with three chains, 20000 iterations, 2000 warmups, and 0.25 thinning. The convergence of the chains is checked using trace plots and histograms. All distributions are Gaussian, and the trace plots demonstrate good convergence and mixing in the algorithm's search and sampling over parameter spaces.

Using the function *add_epred_draws* of the package *tidybayes* (Kay 2020), a series of predictive lines representing 100 draws from the posterior distribution of the Bayesian regression model are overlaid on a scatter plot of observed data (Supplementary Figure 5, top left plots). Predictive intervals around the median of the model predictions are also overlaid on the same scatter plot of observed data (Supplementary Figure 5, top right plots). The predictions show a particularly good fit to observed data for temperatures 28 and 29°C. However, divergence is noticeable at 26 and 27°C, as the survival time data (or lethality buildup rate) are censored at these temperature levels. Consequently, the data points are associated with higher posterior quantiles, where the probability of observing extremely high values is elevated.

Using the function *spread_draws* of the package *tidybayes* (Kay 2020), we extract 10000 random draws from posterior parameter distributions. The parameters of the TTL model encompass the thermal sensitivity parameter $\mathrm{k}$ and both the mean and standard deviation of lethality buildup rates at the reference temperature of 28°C, $\mathrm{mean}(\dot{\mathrm{L}}_\mathrm{r})$ and $\mathrm{SD}(\dot{\mathrm{L}}_\mathrm{r})$, where $\dot{\mathrm{L}}_\mathrm{r}$ is in decadic logarithm scale. Their posterior distributions are all found Gaussian (Supplementary Figure 5, bottom plots). The reference temperature ($\mathrm{T}_\mathrm{r}$) is selected as 28°C which is the lowest of four treatment temperatures that resulted in 100% mussel mortality over the experiment. Nonetheless, the selection of $\mathrm{T}_\mathrm{r}$ do not affect the overall analysis, as any temperature could be selected as the reference point.

**Supplementary Text 5. ABC-SMC approach as implemented using R**

In the *R*-script (refer to https://anonymous.4open.science/r/A-Bayesian-overhaul-of-thermal-tolerance-landscape-models-17F7/), the ABC-SMC approach comprises a series of steps applied separately to each experiment's data and sample names:

1. The script starts by defining a function called *generate_data*. This function creates a set of simulated survival data using the proposed values of the three TTL parameters. It then compares these simulations to our observed survival rates. The discrepancy between the simulations and the observations is quantified using a measurement known as the mean absolute deviation (MAD). Proposal parameters are initial estimates for our model parameters, while the MAD is a simple way of comparing how closely our simulations match the observed data. The *generate_data* function takes parameters *sample_names* (a series of sample names), proposal (proposal values of parameters), *temperature_df* (temperature data frame), *T_ref* (reference temperature), and *observed_surv_df* (observed survival data frame).

   In the process of simulated survival data generation (Monte Carlo simulations), using proposal values of the $\mathrm{mean}(\dot{\mathrm{L}}_\mathrm{r})$ and $\mathrm{SD}(\dot{\mathrm{L}}_\mathrm{r})$ as input for the *rlnorm()* function, a population of 100 $\dot{\mathrm{L}}_\mathrm{r}$ (lethality buildup rates at the reference temperature) is randomly generated assuming a $\log_{10}$-normal distribution. Applying the values of $\dot{\mathrm{L}}_\mathrm{r}$ and the proposal value of $\mathrm{k}$ into Equations 7 and 4 (the equations introduced in the main text), 100 lethality buildup trajectories in response to temperature timeseries were predicted for each experiment. Using Equation 2 and 1, the survival probability trajectory was predicted for the population. Finally, the MAD between the simulated and observed survival probabilities was calculated over all samples separately for the constant heatwave (CHW) and dynamic heatwave (DHW) experiments.

2. The script proceeds to define the *abc_smc* function, which implements the ABC-SMC method. In simpler terms, the ABC-SMC method creates a group of potential solutions, or *particles*, for our problem. Each particle represents a possible set of parameters for our model.

The SMC method starts by creating an initial group of particles (5000 in this case). Each particle represents a possible set of parameters and is formed based on a Gaussian prior distribution. The method then simulates data for each proposed particle using the *generate_data* function (see step 1) and evaluates whether the MAD between the observed and the simulated data for this particle is less than a predetermined threshold, termed the *first epsilon value*. If it meets this criterion, the particle is incorporated into the particle set and assigned a starting weight –probability of resampling the particle at the next iteration – that is equal for all chosen particles at the initial iteration. For our specific application, we used the average and standard deviation of posterior parameter distributions derived from a Bayesian linear regression analysis conducted on data from the CHW experiment (see Supplementary Text 4). It is important to note, though, that the initial values for particles can be chosen somewhat freely and can be further refined through a process of trial and error by utilizing the ABC-SMC algorithm.

The ABC-SMC method goes through a set number of iterations (with a maximum 50 in this case). In each iteration, except the initial iteration, it resamples particles based on their weights, mutates them (i.e., adds some noise), and checks if the mean MAD for the mutated particle's generated data is less than the current epsilon value. If it is, the mutated particle is added to the new particle set. The weights for the new particle set are then calculated based on their proximity to the previous particle set. This is done using a Gaussian kernel, where the variance is given by the *tolerance* parameter. The weights of the chosen particle set are normalized to sum to one, turning the weights into probability values.

Should a particle fail to generate valid data within a maximum number of attempts (10 in this instance), it is omitted from the resampling process. If the success rate (the proportion of particles incorporated into the new particle set) falls below a certain threshold (0.2 in this case), the iterations cease, and the function returns the final particle set, their corresponding weights, the list of mean MAD values, and the iteration count at which the stopping criterion was met. Importantly, this final set of parameter values and their corresponding weights form an approximation of the posterior distribution.

The *abc_smc* function takes the following parameters:

- *sample_names*: This is a vector of identifiers for each sample.
- *n_particles*: The number of particles or proposed parameter sets to use in the SMC algorithm. Using more particles can lead to better coverage of the parameter space but also requires more computational resources.
- *n_iterations*: The number of iterations to run in the SMC algorithm. More iterations can potentially lead to a better fit but also require more computational resources.
- *epsilon_schedule_mean*: A vector of epsilon values for each iteration of the SMC algorithm. Epsilon values determine how closely the simulated data needs to match the observed data for a particle to be accepted into the new particle set. Typically, these values start high and decrease over the iterations.
- *tolerance*: This parameter controls the mutation of particles in the SMC algorithm. It's essentially the standard deviation of the Gaussian noise added to the particles in each iteration. It is specified as a vector with values corresponding to the proposal parameters.

3. Once the *abc_smc* function is defined, the script finally sets up its parameters and runs the ABC-SMC method on the data.

**Supplementary Text 6. Comparison of performance between ABC-SMC and BR**

From the posterior distributions obtained using the Bayesian Regression (BR) or Approximate Bayesian Computation - Sequential Monte Carlo (ABC-SMC) methods, we sample 1000 parameter sets. For each set, the $\mathrm{mean}(\dot{L}_r)$ and $\mathrm{SD}(\dot{L}_r)$ are used as inputs to the *rlnorm()* function to simulate 100 random values of $\dot{L}_r$, for individual mussels, assuming a $\log_{10}$-normal distribution. Utilizing these 100 $\dot{L}_r$ and the corresponding posterior sample of the population rate constant $k$, we predict 100 individual lethality buildup ($L_i$) trajectories under various temperature regimes over time. Subsequently, we derive the population-level survival probability trajectory. Finally, we calculate the Mean Absolute Daily Deviance (MAD) between the simulated survival probabilities and the observed survival probabilities over time for each treatment levels of each experiment. This simulation process and the MAD calculation are iteratively performed for all 1000 sampled posterior parameter sets. Both the simulated and observed survival trajectories, alongside the temperature regimes, are graphically represented in Supplementary Figure 6.

A comparison of the MAD from two parametrization methods, ABC-SMC and BR, across different treatment levels in constant (CHW) and dynamic heatwave (DHW) experiments is presented in Supplementary Figure 7. Treatment levels are denoted by numeric labels (CHW: 26 to 29; DHW: 4 to 5.5). Overall, the ABC-SMC method generally yields lower MAD values, indicating superior model fit, except for the CHW treatment at level 26.

In addition, supplementary Table 2 provides a statistical overview of the two approaches. For CHW, ABC-SMC has a slightly lower median MAD than BR, suggesting it is marginally more precise. The narrower $5^{th}$ to $95^{th}$ percentile range for ABC-SMC indicates more consistent simulation performance. In the DHW setting, ABC-SMC outperforms BR with significantly lower median and mean MAD values and more concentrated distribution, indicating a more robust parametrization method under these conditions. The standard deviation of MAD values is smaller for ABC-SMC in DHW experiments, pointing to less variability compared to BR.

### Supplementary Figures

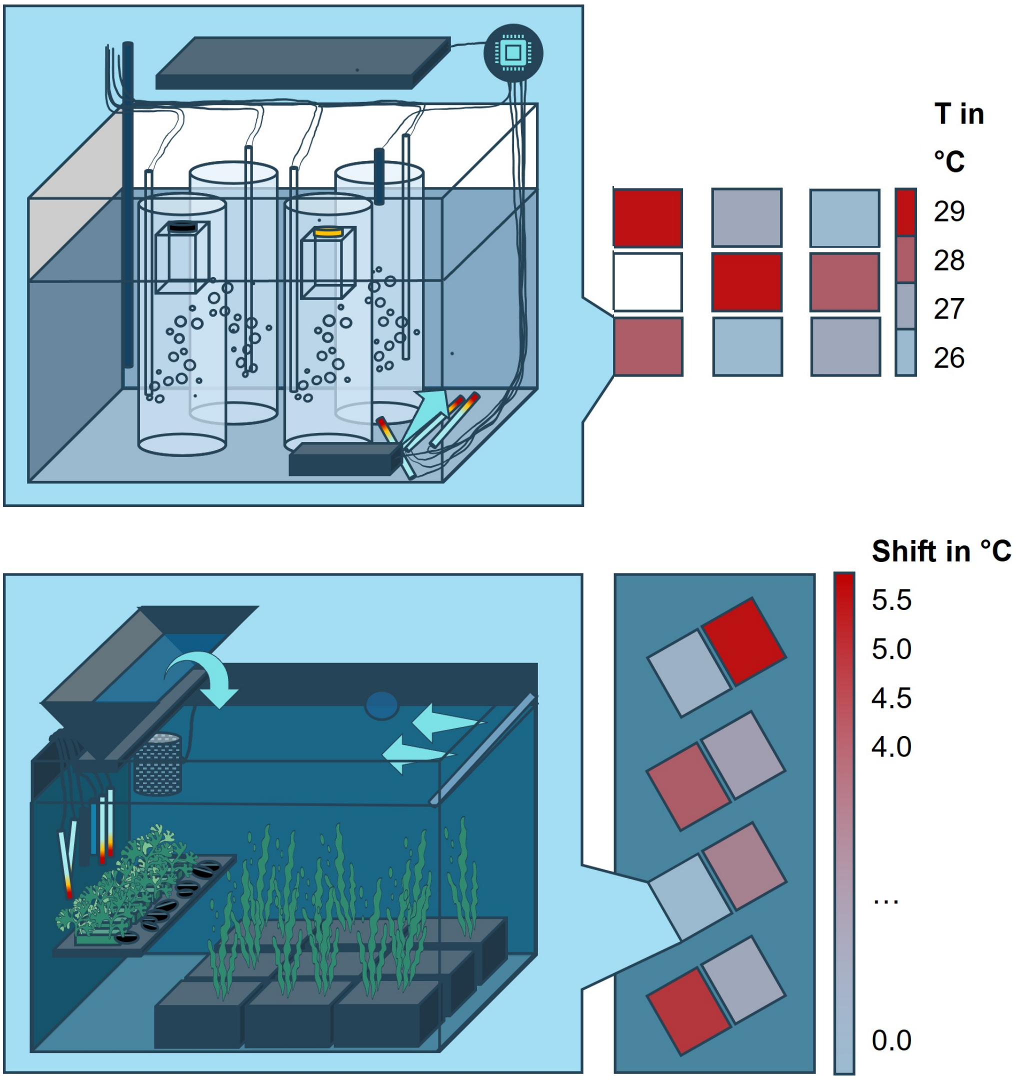


Supplementary Figure 1. Schematic of treatment levels and an exemplary unit of the Kiel Indoor and Outdoor Benthocosm systems (KIBs and KOBs), used in the constant heatwave (CHW) and dynamic heatwave (DHW) experiments, respectively. Each unit of KIBs or KOBs features a computer-controlled heating-cooling system, complemented by a water circulation pump for maintaining consistent water temperatures in the tank. The upper section of the schematic illustrates a KIB unit showcasing the placement of the thermal sensor within one of the standby cylinders, while the two other cylinders accommodate mussels. All cylinders receive an evenly distributed, continuous air supply, regulated via syringes controlling air passage through rigid tubing. The lower section depicts a KOB tank where mussels were housed in PVC cages, alongside regional shallow-water communities including macro-algae *Fucus*, seagrass *Zostera*, and their associated species, all of which underwent identical treatment during the experimental period. Two pumps facilitated water circulation from the tank's depths to its surface and towards a wave generator. The cage containing the mussels was submerged and fastened near the tank's border, adjacent to the wave generator.

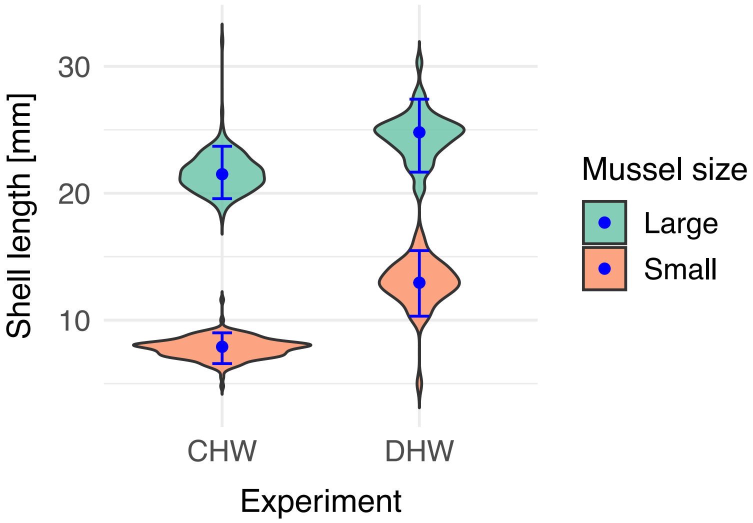


Supplementary Figure 2. Shell length distribution of *Mytilus* mussels of the two size classes at the end of the constant heatwave (CHW) and dynamic heatwave (DHW) experiments.

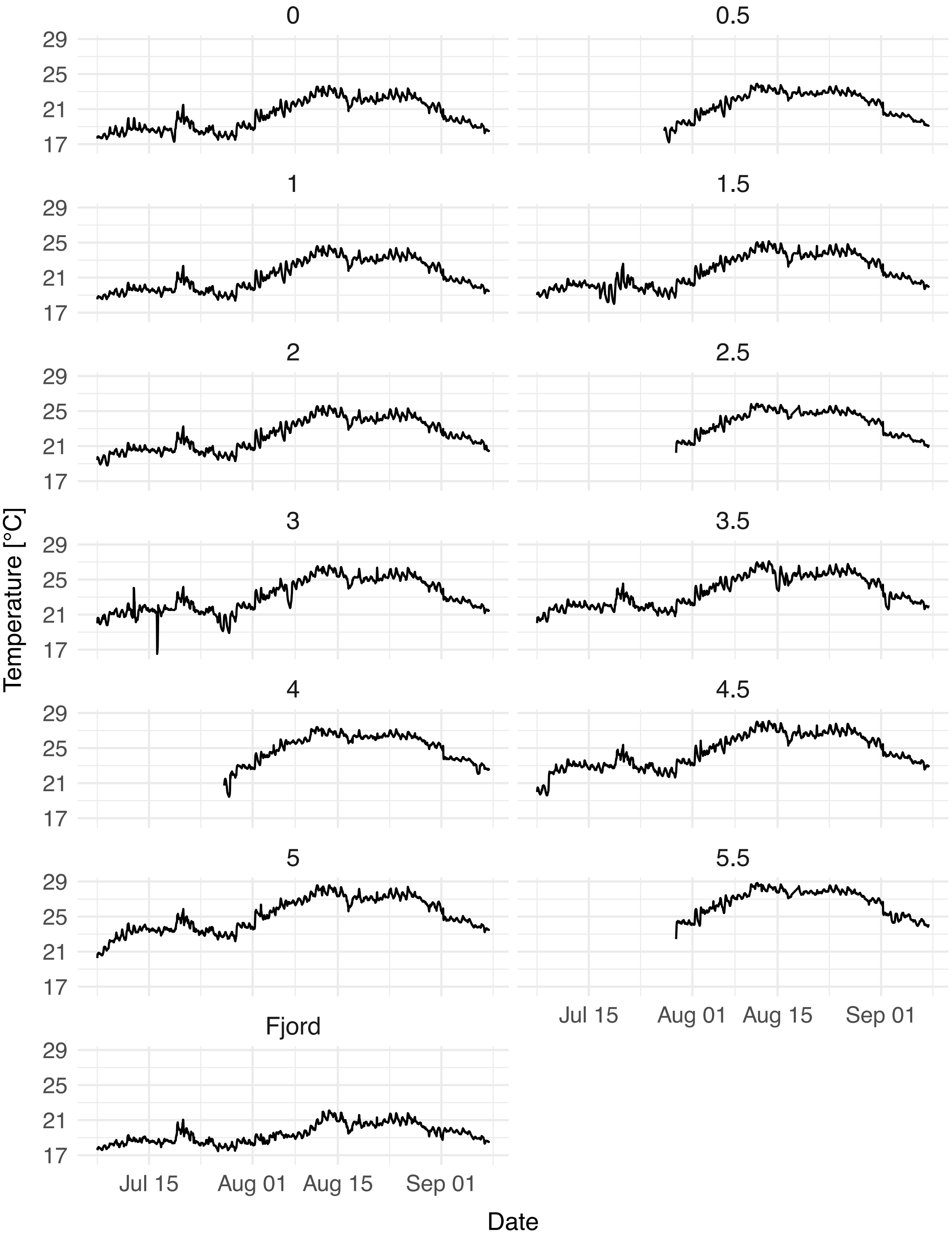


Supplementary Figure 3. Hourly temperature profiles across various treatments (0 to 5.5) during the dynamic heatwave (DHW) experiment, along with natural Kiel fjord conditions, recorded from mid-July to early September. Data were primarily obtained via using GHL sensors—specifically, the platinum resistance thermometer PT1000 (GHL Advanced Technology GmbH, Germany)—installed within the tanks. Subplots represent distinct treatment levels or the fjord conditions during the experimental period. Notably, data gaps are present in treatments 0.5, 2.5, 4, and 5.5, which were compensated by corroborative measurements taken hourly with another set of sensors (refer to Supplementary Figure 4).

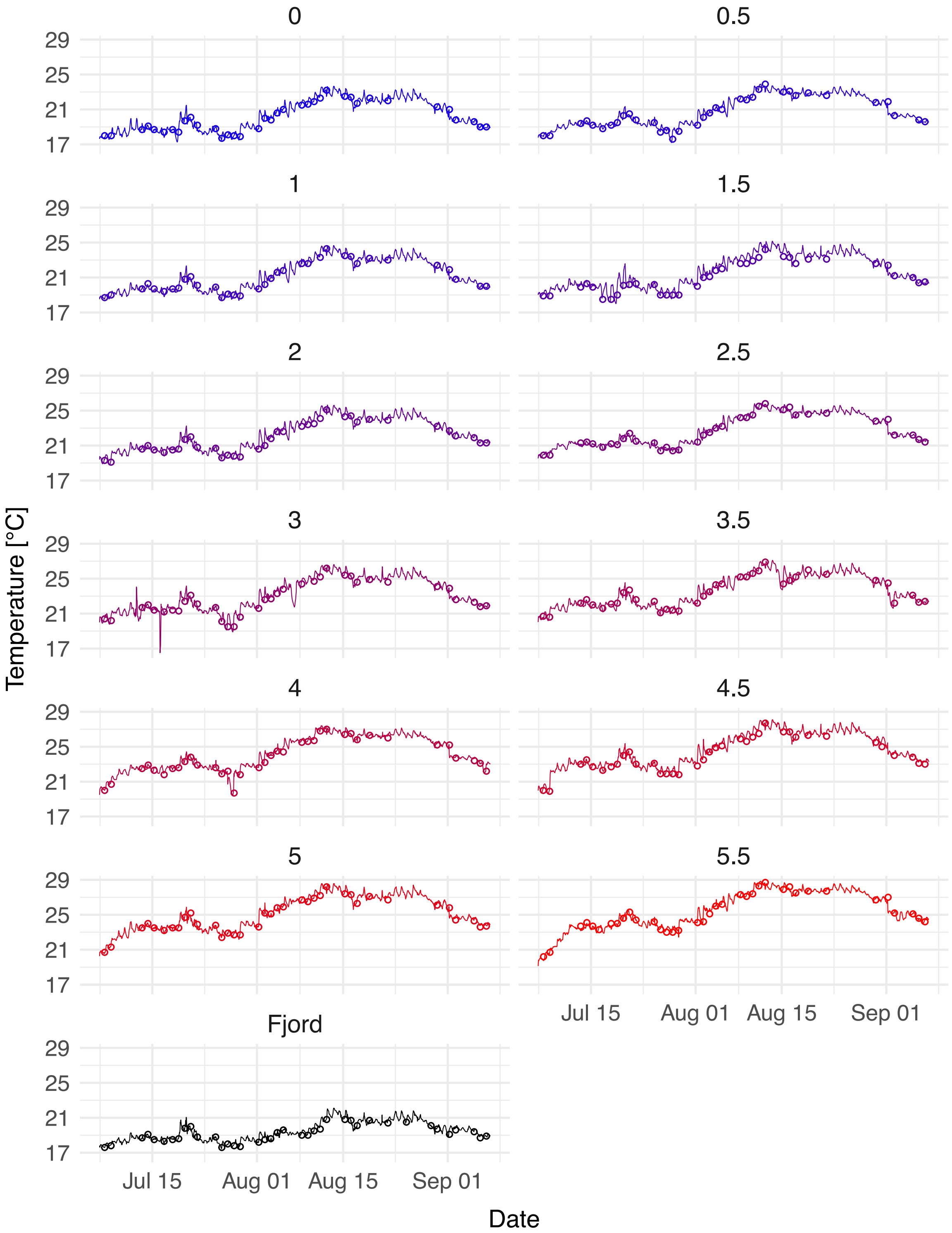


Supplementary Figure 4. Association of temperature datasets for all experimental treatments alongside fjord conditions, showing GHL sensor recordings (lines) supplemented with hourly data collected by Hach sensor (Dissolved Oxygen Sensor, Hach Lange GmbH, Germany), for missing intervals, against corroborative WTW sensor measurements (circles). Readings of WTW sensor (Xylem Analytics GmbH, Germany), taken approximately daily, confirm the precision of hourly GHL data. Each subplot tracks temperature for a specific treatment or the fjord, validating the temperature control and monitoring system's efficacy throughout the experiment.

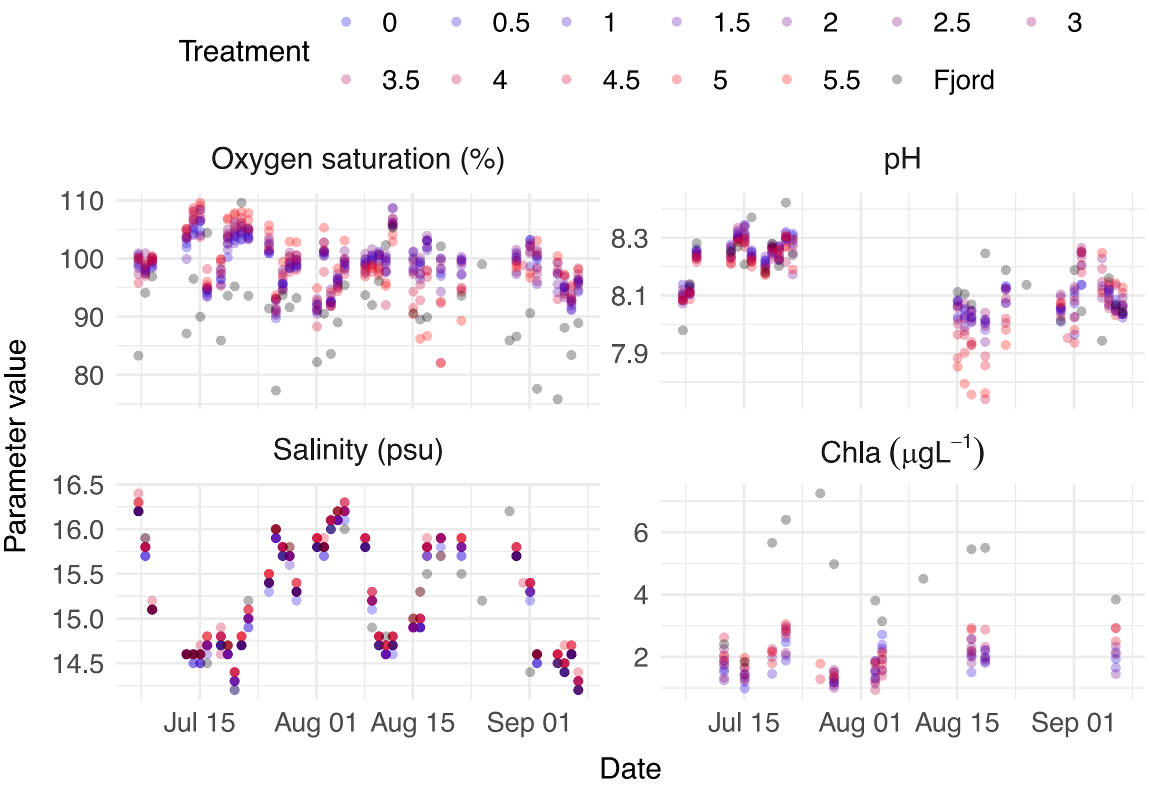


Supplementary Figure 5. Daily recorded values of oxygen saturation, pH, and salinity, along with weekly measurements of chlorophyll-a concentrations, across all treatment levels of the dynamic heatwave (DHW) experiment, and fjord conditions. Oxygen saturation, pH, and salinity were recorded daily using a portable multiparameter device (WTW Multi 3630 IDS with pH Sentix 940 ID, Salinity TetraCon 925 IDS and Oxygen FDO 925 IDS sensors). Chlorophyll concentrations were assessed weekly with Cyclops 7f Chl sensors (Turner Designs, USA). These parameters were monitored to confirm that their values remained within a range that would not be extreme to the mussels, independent of the temperature effects.

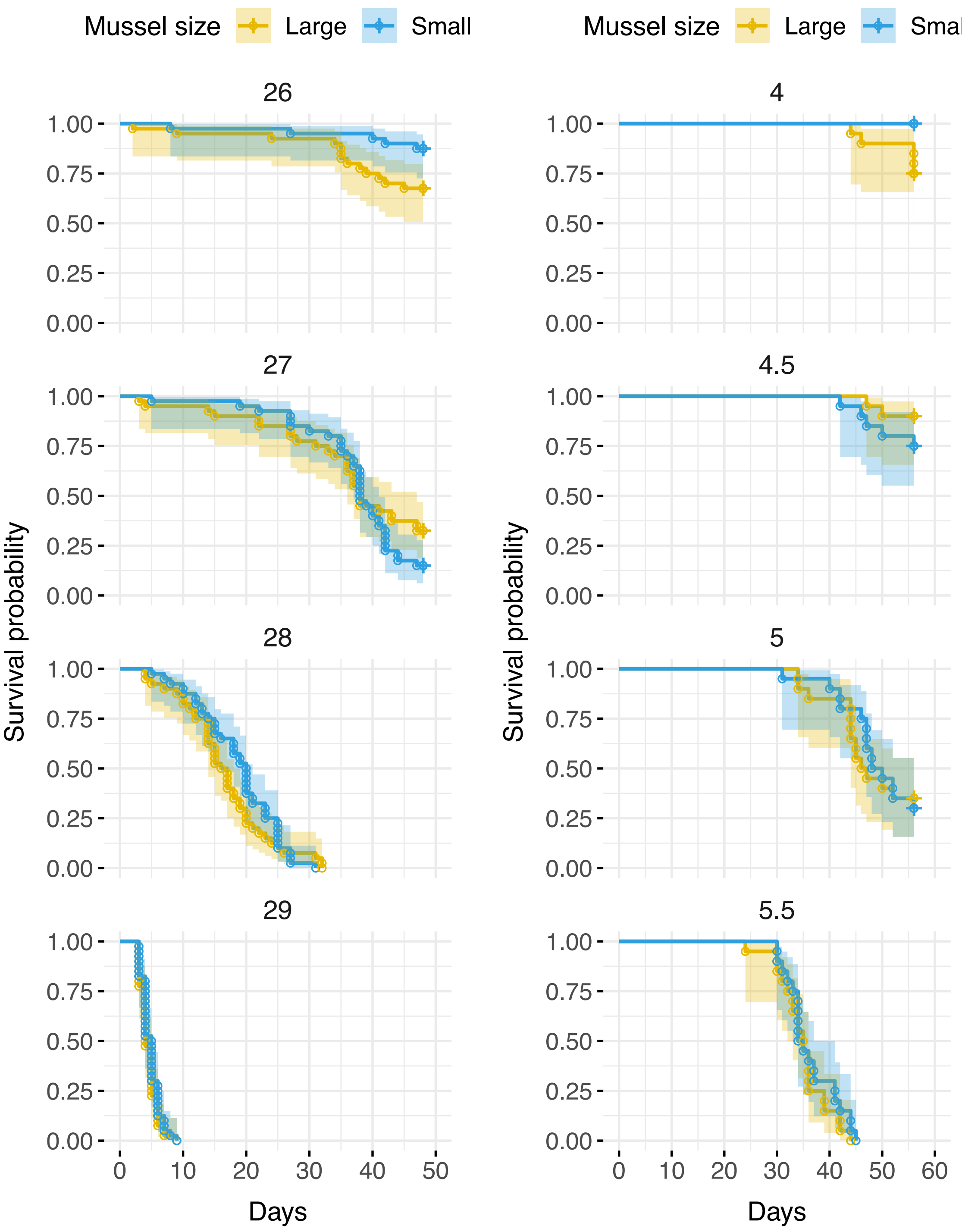


Supplementary Figure 6. Survival trajectories of large and small *Mytilus* mussels under the treatments reflecting both constant (CHW) and dynamic heatwave (DHW) conditions, designated by the identifiers 26, 27, 28, 29 for CHW, and 4, 4.5, 5, 5.5 for DHW, respectively. Survival probabilities are plotted as discrete data points (circles), with interpolated survival curves derived from Kaplan-Meier estimations overlaid as solid lines. Shaded areas represent 95% confidence intervals, illustrating the variance within each size class across treatment levels.

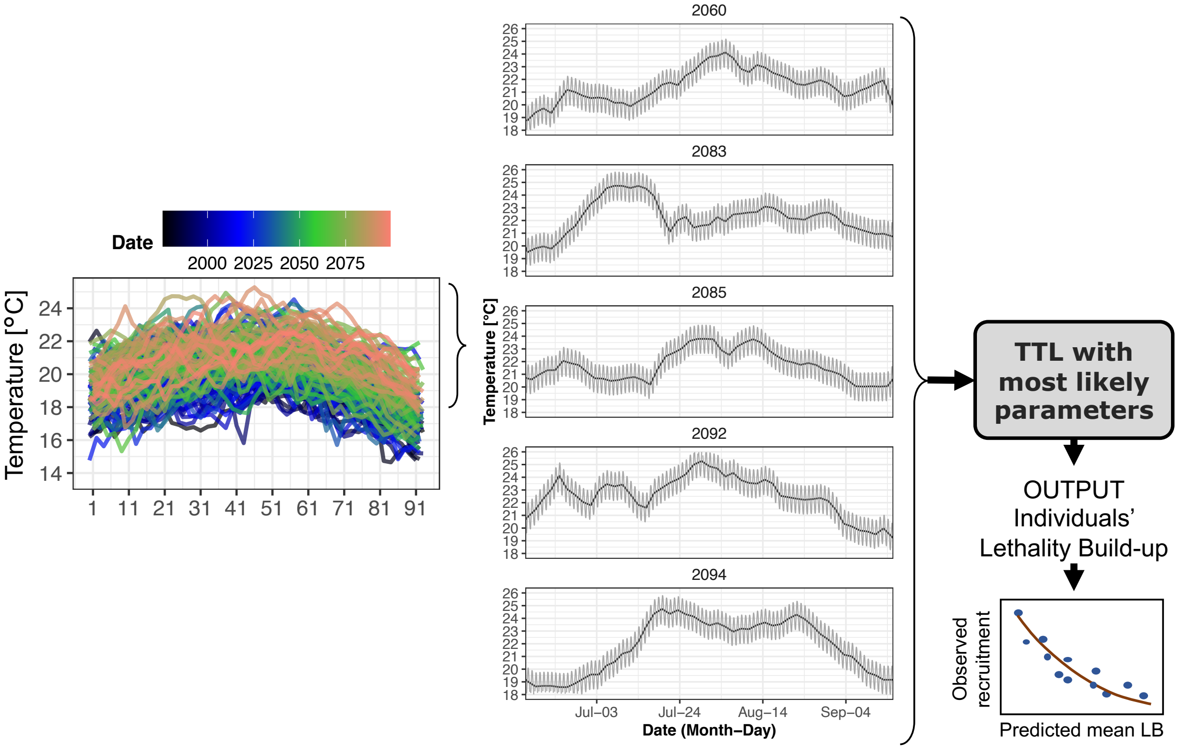


Supplementary Figure 7. Projections of summer temperatures for the current century at the Kiel Fjord entrance site under the Representative Concentration Pathway 8.5 scenario were refined to an hourly scale (left side plot). The five warmest temperature regimes, both with and without a representative daily temperature fluctuation cycle (middle plots), were input into the TTL using the most likely parameter set determined by the ABC-SMC approach. This allowed for predictions of individual lethality buildup trajectories and population survival across each summer (not depicted here). The predicted average lethality buildup served as an input for the established (non-causal) relationship (as shown in Figure 5 of the main text) to forecast recruitment probabilities under the different summer scenarios.

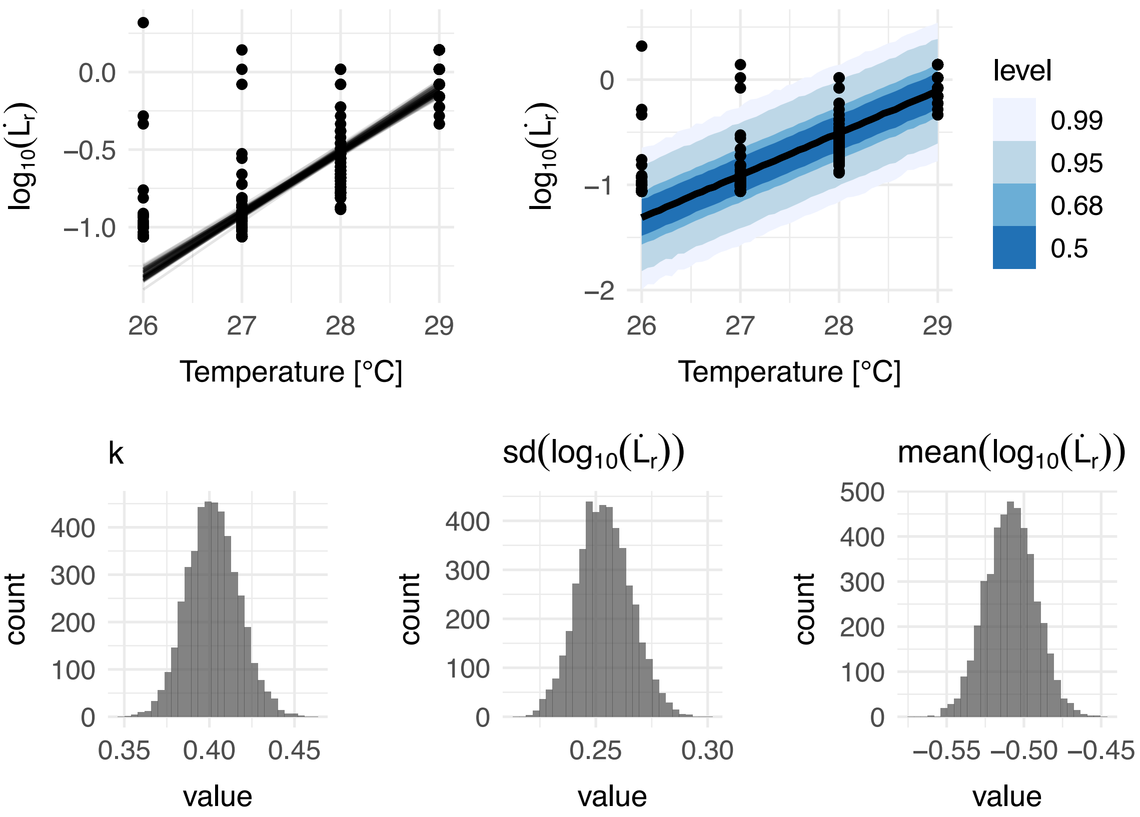


Supplementary Figure 8. Posterior predictions of the Bayesian log-linear regression modelling of the lethality buildup rate ($\dot{L}$; at log10 scale) over constant temperatures and the observations (dots). The predicted lines of best fits (mean responses) are shown as black lines (top left). The predicted quantile ranges of the response are presented in various shades of blue around the predicted best-fit line (top right). Posterior distributions of the TTL parameters, including the thermal sensitivity parameter ($k$), and the mean and standard deviation of the decadic logarithm of the lethality buildup rate at the reference temperature are presented as histograms (bottom left to right).

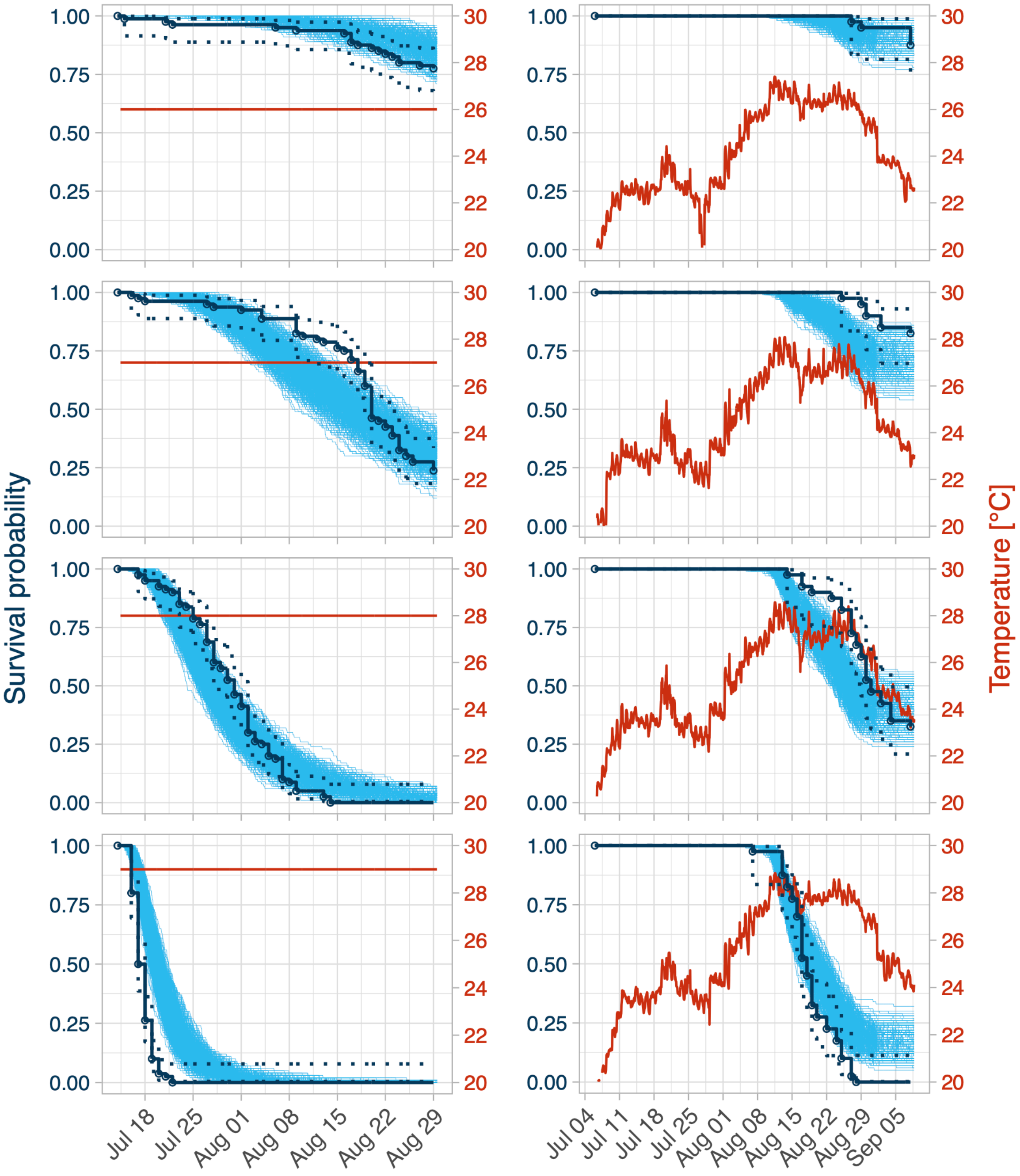


Supplementary Figure 9. Monte Carlo simulation of mussel survival probabilities at constant or dynamic heatwave regimes (left versus right-side columns) based on the whole posterior parameter sets from the Bayesian regression (see Supplementary text 3). Simulated trajectories are presented as blue lines. The observed survival probabilities are presented as black circles. The corresponding Kepler-Meyer predictions are interpolated (continuous lines) with 95% confidence interval (dotted lines). Thermal regimes are displayed in red.

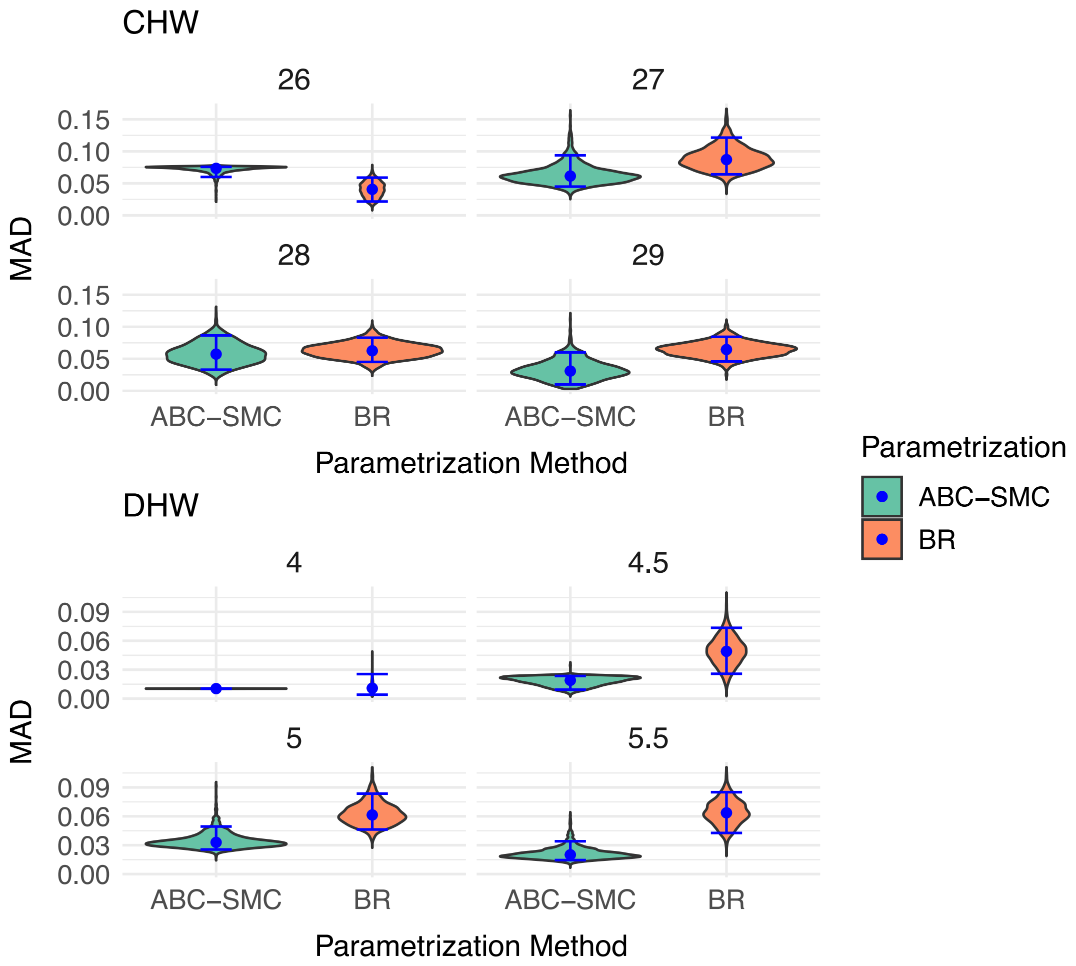


Supplementary Figure 10. Distributions of the mean absolute deviance (MAD) for constant heatwave (CHW) and dynamic heatwave (DHW) experiments across various treatment levels. Violin plots display the MAD values for both ABC-SMC and BR parametrization methods, with blue diamonds representing the median and blue bars showing the $5^{th}$ to $95^{th}$ percentile range. Treatment levels are displayed within each facet.

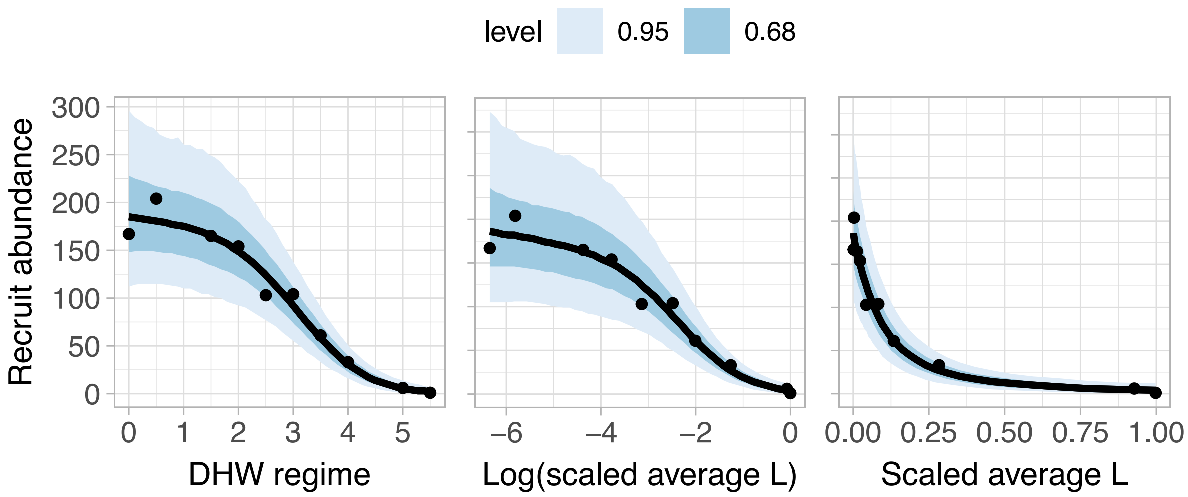


Supplementary Figure 11. Patterns of mussel recruit abundance in relation to dynamic heatwave (DHW) regimes and average lethality buildup predicted by ABC-SMC. Left panel: Observed logistic decline in recruit abundance with increasing DHW regime. Middle panel: Non-causal correlation between recruit abundance and the logarithm of scaled average lethality, showing a similar declining trend. Right panel: Non-causal correlation between recruit abundance and non-logarithmic scaled average lethality. Data points correspond to the observed recruitment values, and shaded areas denote the 95% (light blue) and 68% (dark blue) credible intervals around the median predicted using the Bayesian regression model.

### Supplementary Tables

Supplementary Table 1. Cox proportional hazards model analysis detailing the effects of size class and temperature treatment on *Mytilus* mussel survival across constant (CHW) and dynamic (DHW) heatwave experiments. Size classes included small and large (with large serving as the control), and temperature treatments were set at 26°C (control), 27°C, 28°C, and 29°C for CHW, and 4 (control), 4.5, 5, and 5.5 for DHW. Displayed are the model's coefficients, standard errors, test statistics, and 95% confidence intervals. Crucially, size class effects were not statistically significant in either CHW (HR=0.87, p>0.05) or DHW (HR=0.80, p>0.05) settings, supporting the pooled analysis of size classes for advanced ABC-SMC procedures. Temperature treatments, however, were significant predictors of survival, particularly at higher temperature levels, underscoring the thermal sensitivity of mussel populations.

<table>
<tr><th>Experiment</th><th>Non-control levels</th><th>coef</th><th>exp(coef)</th><th>se(coef)</th><th>z</th><th>Pr(>|z|)</th><th>2.5 %</th><th>97.5 %</th></tr>
<tr><td rowspan="4">CHW</td><td>Small</td><td>-0.14</td><td>0.87</td><td>0.13</td><td>-1.10</td><td>0.27</td><td>-0.40</td><td>0.11</td></tr>
<tr><td>27</td><td>1.63</td><td>5.08</td><td>0.27</td><td>6.01</td><td>0.00</td><td>1.10</td><td>2.16</td></tr>
<tr><td>28</td><td>4.32</td><td>75.45</td><td>0.35</td><td>12.47</td><td>0.00</td><td>3.64</td><td>5.00</td></tr>
<tr><td>29</td><td>7.58</td><td>1952.45</td><td>0.47</td><td>16.23</td><td>0.00</td><td>6.66</td><td>8.49</td></tr>
<tr><td rowspan="4">DHW</td><td>Small</td><td>-0.22</td><td>0.80</td><td>0.23</td><td>-0.98</td><td>0.33</td><td>-0.67</td><td>0.22</td></tr>
<tr><td>4.5</td><td>0.37</td><td>1.45</td><td>0.59</td><td>0.63</td><td>0.53</td><td>-0.78</td><td>1.52</td></tr>
<tr><td>5</td><td>2.19</td><td>8.97</td><td>0.49</td><td>4.48</td><td>0.00</td><td>1.23</td><td>3.15</td></tr>
<tr><td>5.5</td><td>4.81</td><td>122.25</td><td>0.55</td><td>8.66</td><td>0.00</td><td>3.72</td><td>5.89</td></tr>
</table>

Supplementary Table 2. Summarized temperature regimes applied in the 64-day dynamic heatwave (DHW) experiment are characterized by various temperature statistics, including the median, 5$^{th}$ and 95$^{th}$ percentiles, mean, standard deviation (SD), minimum (min), and maximum (max) temperatures [°C]. 'Fjord' denotes the natural temperature regime in the Kiel fjord. The treatments labeled from 0 to 5.5 correspond to different warming scenarios in the DHW experiment, with each level representing an incremental increase in thermal intensity. This dataset provides insights into the specific thermal conditions under which the survival of mussels was examined.

| Treatment | Duration (d) | Median | P5 | P95 | mean | SD | min | max |
|---|---|---|---|---|---|---|---|---|
| Fjord | 64 | 19.3 | 17.9 | 21.3 | 19.4 | 1.1 | 17.4 | 22.1 |
| 0 | 64 | 19.9 | 17.9 | 23.0 | 20.3 | 1.7 | 17.3 | 23.6 |
| 0.5 | 64 | 20.4 | 18.5 | 23.4 | 20.8 | 1.7 | 17.2 | 23.9 |
| 1 | 64 | 20.8 | 18.9 | 24.0 | 21.2 | 1.8 | 18.4 | 24.7 |
| 1.5 | 64 | 21.3 | 19.1 | 24.5 | 21.7 | 1.8 | 18.0 | 25.1 |
| 2 | 64 | 22.0 | 19.8 | 25.0 | 22.3 | 1.8 | 18.8 | 25.7 |
| 2.5 | 64 | 22.3 | 20.5 | 25.4 | 22.8 | 1.8 | 19.5 | 25.8 |
| 3 | 64 | 22.8 | 20.5 | 26.0 | 23.2 | 1.9 | 16.5 | 26.6 |
| 3.5 | 64 | 23.3 | 21.2 | 26.3 | 23.6 | 1.8 | 20.0 | 27.1 |
| 4 | 64 | 23.8 | 21.3 | 26.9 | 24.2 | 1.9 | 19.4 | 27.4 |
| 4.5 | 64 | 24.3 | 22.0 | 27.5 | 24.6 | 1.9 | 19.6 | 28.1 |
| 5 | 64 | 24.8 | 22.4 | 28.0 | 25.1 | 1.9 | 20.3 | 28.6 |
| 5.5 | 64 | 25.2 | 22.0 | 28.4 | 25.5 | 2.1 | 19.1 | 28.8 |

Supplementary Table 3. Comparison of the mean absolute deviance (MAD) by the parametrization method. The table compares MAD values obtained using ABC-SMC and BR methods, illustrating median and percentile range (5% and 95%) for constant (CHW) and dynamic (DHW) heatwave experiments. The metrics allow for an assessment of accuracy and consistency of each parametrization method across different experimental conditions, with lower values signifying more precise predictions.

| Parametrization | Experiment | Median MAD | P5 | P95 |
|---|---|---|---|---|
| ABC-SMC | CHW | 0. 0572 | 0. 0340 | 0. 0758 |
| ABC-SMC | DHW | 0. 0203 | 0. 0104 | 0. 0371 |
| BR | CHW | 0.0641 | 0.0550 | 0.0744 |
| BR | DHW | 0.0469 | 0.0370 | 0.0577 |

Supplementary Table 4. Summary statistics of Thermal Tolerance Landscape (TTL) model parameters estimated by an Approximate Bayesian Computation-Sequential Monte Carlo (ABC-SMC) approach for two experimental conditions: constant heatwave (CHW) and dynamic heatwave (DHW) regimes. The summary includes the median and 5th-95th percentile range for $mean(\dot{L_r})$, $SD(\dot{L_r})$, and k, where $\dot{L_r}$ is in $\log_{10}$ or original scale.

| Parameter | Experiment | Median (Log10) | P5 (Log10) | P95 (Log10) | Median (original) | P5 (original) | P95 (original) |
|---|---|---|---|---|---|---|---|
| k | CHW | 0.482 | 0.410 | 0.548 | 3.032 | 2.573 | 3.536 |
| | DHW | 0.535 | 0.454 | 0.618 | 3.430 | 2.844 | 4.152 |
| $\mathrm{mean}(\dot{L}_r)$ | CHW | -0.500 | -0.543 | -0.448 | 0.316 | 0.287 | 0.356 |
| | DHW | -0.472 | -0.504 | -0.445 | 0.337 | 0.313 | 0.359 |
| $\mathrm{SD}(\dot{L}_r)$ | CHW | 0.195 | 0.154 | 0.241 | 0.383 | 0.322 | 0.452 |
| | DHW | 0.131 | 0.103 | 0.165 | 0.307 | 0.262 | 0.358 |
